# ZBTB20 is Essential for Cochlear Maturation and Hearing in Mice

**DOI:** 10.1101/2022.12.23.521726

**Authors:** Zhifang Xie, Xian-Hua Ma, Qiu-Fang Bai, Jie Tang, Jian-He Sun, Fei Jiang, Wei Guo, Chen-Ma Wang, Rui Yang, Yin-Chuan Wen, Fang-Yuan Wang, Yu-Xia Chen, Hai Zhang, David Z. He, Matthew W. Kelley, Shiming Yang, Weiping J. Zhang

## Abstract

The mammalian cochlear epithelium undergoes substantial remodeling and maturation before the onset of hearing. However, very little is known about the transcriptional network governing cochlear late-stage maturation and particularly the differentiation of its lateral non-sensory region. Here we establish ZBTB20 as an essential transcription factor required for cochlear terminal differentiation and maturation and hearing. ZBTB20 is abundantly expressed in the developing and mature cochlear non-sensory epithelial cells, with transient expression in immature hair cells and spiral ganglion neurons. Otocyst-specific deletion of Zbtb20 causes profound deafness with reduced endolymph potential in mice. The subtypes of cochlear epithelial cells are normally generated but their postnatal development is arrested in the absence of ZBTB20, as manifested by an immature appearance of the organ of Corti, malformation of tectorial membrane, a flattened spiral prominence, and a lack of identifiable Boettcher cells. Furthermore, these defects are related with a failure in the terminal differentiation of the non-sensory epithelium covering the outer border Claudius cells, outer sulcus root cells and spiral prominence epithelial cells. Transcriptome analysis shows ZBTB20 regulates genes coding for tectorial membrane proteins in the greater epithelial ridge, and those preferentially expressed in root cells and spiral prominence epithelium. Our results point to ZBTB20 as an essential regulator for postnatal cochlear maturation and particularly for the terminal differentiation of cochlear lateral non-sensory domain.

## Introduction

The sensory epithelium, the organ of Corti, within the mammalian cochlea is a complex structure consisting of mechanosensory hair cells (HCs) and specialized non-sensory supporting cells (SCs). In mice, the cochlear epithelium is immature at birth and continues to remodel and mature structurally and functionally until it reaches an adult-like appearance at around P21 (1,2). The morphologic maturation process is characterized by the regression of the greater epithelial ridge (GER) leading to the formation of the inner sulcus, the detachment of the tectorial membrane (TM) from the organ of Corti, the opening of the tunnel of Corti, the formation of the spaces of Nuel, and dramatic changes in the shape and size of the outer border cells including Hensen cells, Claudius cells, and Boettcher cells(3).

Another morphologic change during this period is the development of root cells in the outer sulcus region(2). The outer sulcus epithelia initially present as a simple layer of cuboidal-shaped cells bordering the scala media, but shortly after birth some of these cells infiltrate into the spiral ligament and go on to differentiate as distinct root cells with extensive branched processes extending deep into the spiral ligament. At the same time, the surfaces of the root cell bodies become covered by Claudius cells and spiral prominence (SP) epithelial cells and are no longer directly in contact with the endolymph(2, 4, 5). Simultaneously, the SP becomes identifiable as a triangular structure located between the lower margin of the stria vascularis (SV) and the upper border of the outer sulcus(6).

Paralleling the morphologic maturation of the cochlea, the ionic composition of the endolymph matures leading to a high K^+^ and low Na^+^, followed by the generation of an endocochlear potential (EP) of approximately +80 mV relative to perilymph (7-9). The positive EP serves as the main driving force for mechanoelectrical transduction by HCs, the deterioration of which leads to hearing loss or deafness (10). In comparison with the early-stages of cochlear specification and differentiation(11-13), relatively little is known about the transcriptional networks governing the late-stages differentiation and terminal maturation of cochlear epithelium, particularly the differentiation of outer border Claudius and Boettcher cells, outer sulcus root cells and the morphogenesis of the SP.

Zinc finger protein Zbtb20 is a member of the broad complex tramtrack bric-a-brac (BTB) zinc-finger (ZnF) family (14, 15). We and others have reported that Zbtb20 functions primarily as a transcription factor and plays essential roles in the development of multiple organ systems (16-21). Interestingly, we noticed that *Zbtb20* null mice display head tilt and circling behavior, suggestive of an inner ear defect. In humans, de novo heterozygous missense variants in *ZBTB20* (MIM* 606025) causes Primrose syndrome (MIM# 259050), which is a rare genetic condition characterized by cognitive deficits, unusual facial features, and calcified external ears (22, 23). Strikingly, affected individuals frequently exhibit sensorineural hearing loss in early childhood (23), suggesting ZBTB20 may be required for auditory development. However, its cellular and molecular mechanism remains an enigma. Here, we show that ZBTB20 is abundantly and persistently expressed by various cochlear non-sensory epithelial cells, including supporting cells, outer border cells, outer sulcus root cells and SP epithelial cells. Conditional knockout of *Zbtb20* in otic vesicle results in profound deafness with markedly reduced EP and permanently arrested postnatal development of cochlear epithelium covering the outer border, outer sulcus and SP regions. Taken together, our findings establish ZBTB20 as an essential regulator in postnatal cochlear maturation.

## Results

### Global knockout of *Zbtb20* causes deafness

Global *Zbtb20* knockout mice (*Zbtb20*^-/-^) displayed head tilt and circling behavior, suggesting an inner ear defect(16). To assess their hearing function, we performed auditory brainstem response (ABR) tests at postnatal day 21 (P21), when the auditory system becomes mature. *Zbtb20*^*-/-*^ mice displayed profound hearing loss at all frequencies tested (*SI Appendix*, Fig. S1). As *Zbtb20*^*-/-*^ mice exhibit high premature mortality, severe hypoglycemia, and developmental defects in multiple organs (e.g. brain, pituitary, and bone) (16, 18, 21, 24), it was complicated and hard to explore the mechanism about the hearing phenotype.

### ZBTB20 is differentially expressed in cochlear epithelial cells

To elucidate the role of *Zbtb20* in hearing, we first examined the spatiotemporal expression pattern of ZBTB20 in murine cochleae at different developmental stages by immunostaining. An initial expression of ZBTB20 protein was detected in the lesser epithelial ridge (LER) of cochlear duct at embryonic day 13.5 (E13.5), with poor expression in the GER (*SI Appendix*, Fig. S2*A*). This expression pattern was maintained at E16.5, when ZBTB20 was expressed strongly in Myosin7a-positive HCs and the adjacent SCs, as well as epithelial cells in the outer sulcus, SV and Reissner’s membrane (RM), but undetectable in the GER (Fig. 1*A*).

**Figure 1.**
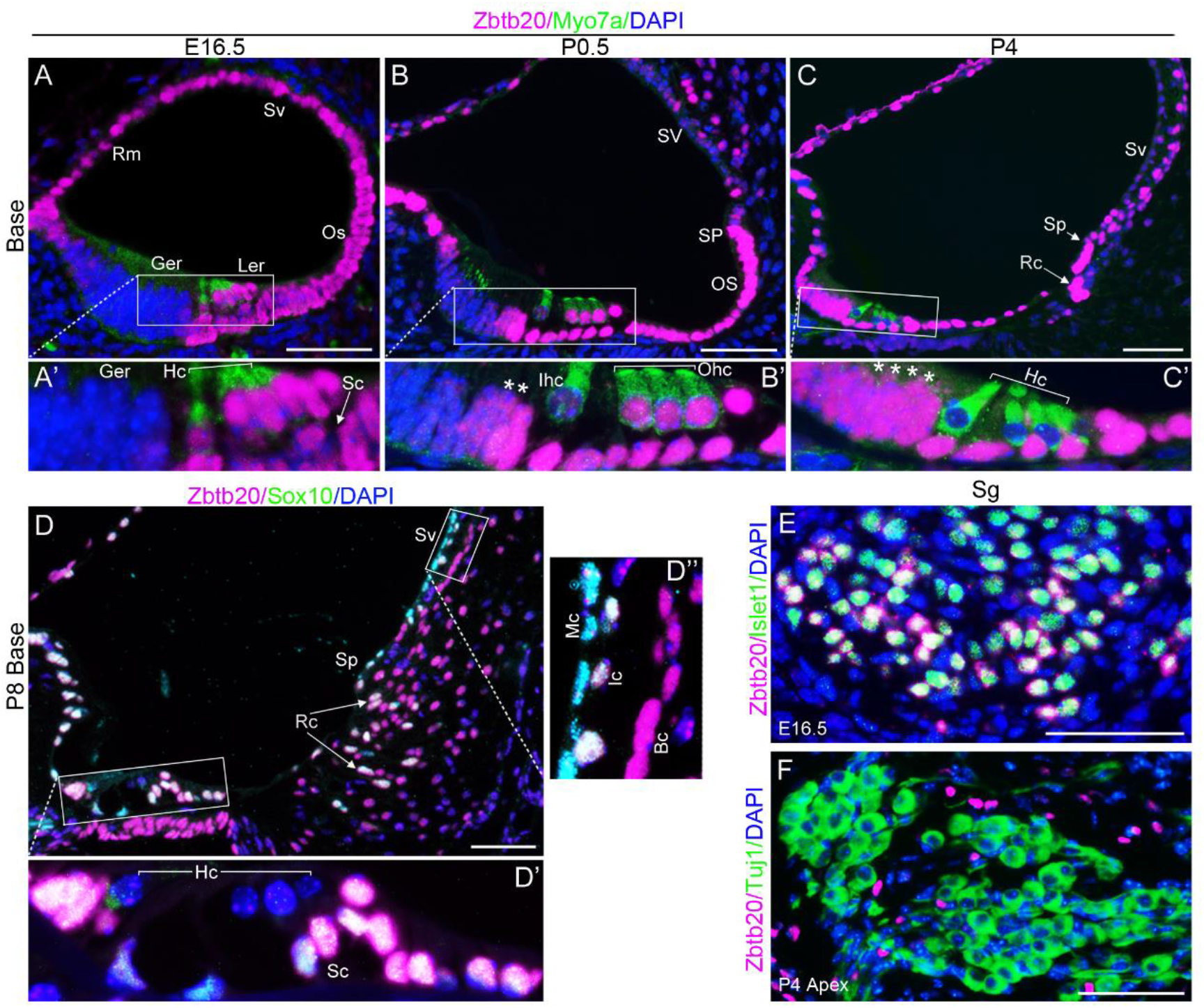
ZBTB20 is differentially expressed in cochlear epithelial cells. (**A-D**) Double-immunofluorescence staining showing spatial and temporal expression pattern of ZBTB20 (magenta) in cochlea. (**A**) ZBTB20 was expressed strongly in Myosin7a-positive (green) hair cells (Hc) and the adjacent supporting cells (Sc), outer sulcus (Os), stria vascularis (Sv), Reissner’s membrane (Rm) but undetectable in greater epithelial ridge (Ger) at E16.5. (**B-C**) ZBTB20 staining intensity becomes weaker in Myosin7a-positive inner hair cells (Ihc) at P0.5 (**B**) and undetectable in all HCs at P4 (**C**), while increased in the lateral and throughout GER at P0.5 and P4, respectively (indicated with *). Note the developing root cells (Rc) and spiral prominence epithelium (Sp) showing strong signals for ZBTB20. Idc, interdental cells, Ohc, outer hair cells. (**D**) At P8, ZBTB20 was strongly expressed by cochlear Sox10-positive (green) non-sensory epithelial cells including supporting cells (Sc) and root cells (Rc, indicated by arrowheads), with the exception of Sox10-positive SV marginal cells (Mc), which showed very weak staining. Ic and Bc indicate SV intermediate cell and basal cell, respectively; Ler, lesser epithelial ridge. (**A**’, **B**’, **C**’, **D**’ and **D**’’) are magnified views of indicated boxed areas in (**A**-**D**). (**E and F**) Double-immunofluorescence staining showing ZBTB20 (magenta) was expressed by a subset of Islet-positive (green) spiral ganglion (Sg) neurons at E16.5 but undetectable in Tuj1-positive (green) SG neurons at P4. Nuclei were stained with DAPI (blue). n=4 mice for each developmental stage. Scale bars in all panels=50 µm. See also *SI Appendix* Fig. S2 and S3.

Shortly after birth (P0.5∼P4), ZBTB20 protein diminished in Myosin7a-positive inner hair cells (IHCs) and then outer hair cells (OHCs), a pattern that parallels the progression of HC maturation from IHCs to OHCs (25, 26). Simultaneously, ZBTB20 expression increased in the GER in a basal-to-apical gradient (Fig. 1*B-C*). This pattern is consistent with the wave of cellular remodeling that occurs in the GER (27). Intense expression signals for ZBTB20 were observed in SCs, outer sulcus/root cells and SP epithelium, and moderate signals in the RM and SV at both P0.5 and P4 (Fig. 1*B-C*). This expression pattern was also confirmed by double immunostaining for ZBTB20 and Sox10 (*SI Appendix*, Fig. S 2*B-C*), which is expressed in the nuclei of all SCs and other cochlear non-sensory epithelia facing the endolymph, but not in HCs (28). At P8, ZBTB20 was strongly expressed by all cochlear Sox10-positive non-sensory epithelial cells except for SV marginal cells, which displayed very weak staining signal (Fig. 1*D*). Of note, in the outer sulcus region, the developing Sox10-positive root cells showed strong expression signals of ZBTB20. This expression pattern of ZBTB20 was maintained at P10 and throughout adulthood (*SI Appendix* Fig. S2*D-E*).

In the spiral ganglions (SG) regions, weak staining for ZBTB20 was initially detected at E13.5 (*SI Appendix* Fig. S2*A*). At E16.5, when SG neurons and glia can be identified using different markers, ZBTB20 was detected in a subset of Islet1-positive neurons (Fig. 1*E*). Expression of ZBTB20 was preserved in a subset of TuJ1 (β-tubulin III, a pan neuronal marker)-positive neurons in the most apical and lower basal but not the middle SGs at P0.5 (*SI Appendix* Fig. S3*A-B*), and became undetectable in Tuj1-positive neurons in all turns at P4 and thereafter (Fig. 1*F* and *SI Appendix* Fig. S3*C*). Co-localization of ZBTB20 with Sox10 (a pan marker for SG glial cells) was undetectable in SG regions at E16.5, P0.5, or P4 (*SI Appendix* Fig. S3*D-F*), suggesting ZBTB20 is not expressed by SG glial cells. ZBTB20 was also detected in some scattered cells expressing neither Tuj1 nor Sox10 (likely mesenchyme cells) at P0.5 (*SI Appendix* Fig. S3*B-E*) and thereafter (Fig.1*F, SI Appendix* Fig. S3*C, F*).

In summary, ZBTB20 is abundantly expressed in the developing and mature cochlear non-sensory epithelial cells, with transient expression in immature HCs and a subset of SG neurons.

### Conditional knockout of *Zbtb20* causes EP reduction and deafness

We then generated otic vesicle-specific *Zbtb20* knockout mice (hereafter referred to as OV-ZB20KO) to determine the role of *Zbtb20* in cochlear development. To this aim, we crossed *Zbtb20-floxed* mice (*Zbtb20*^*flox*^) with *Foxg1-Cre* transgenic mice, a line that has been shown to mediate recombination specifically in the otic vesicle by E8.5 (29). Immunofluorescence analysis confirmed the absence of ZBTB20 expression in OV-ZB20KO cochlear ducts and SG at E13.5 (*SI Appendix* Fig. S4*A-B*), the earliest stage at which ZBTB20 expression was detected in wild-type (WT) counterparts. Consistently, expression of ZBTB20 was observed in control but not OV-ZB20KO sensory and non-sensory epithelia surrounding the scala media at P0.5 (*SI Appendix* Fig. S4*C-F*) or P8 (Fig. 2*A-B*), including Myosin7a-positive HCs at P0.5 (*SI Appendix* Fig. S4*C-D*), Sox10-positive SCs, outer sulcus/root cells, epithelial cells in the GER, as well as marginal cells in the SV (P0.5) (Fig. 2*A-B*, *SI Appendix* Fig. S4*C-F*). In SG, expression of ZBTB20 was detectable in a subset of Tuj1-positive neurons from control but not from OV-ZB20KO pups at P0.5 (*SI Appendix* Fig. S4*G-H*). Of note, the tympanic border cells beneath the basilar membrane and fibrocytes in the lateral wall, both of which are of non-OV origin, displayed comparable ZBTB20 expression between the two genotypes at P8 (Fig. 2*A-B*), confirming the specific deletion of *Zbtb20* in OV-derived cells.

**Figure 2.**
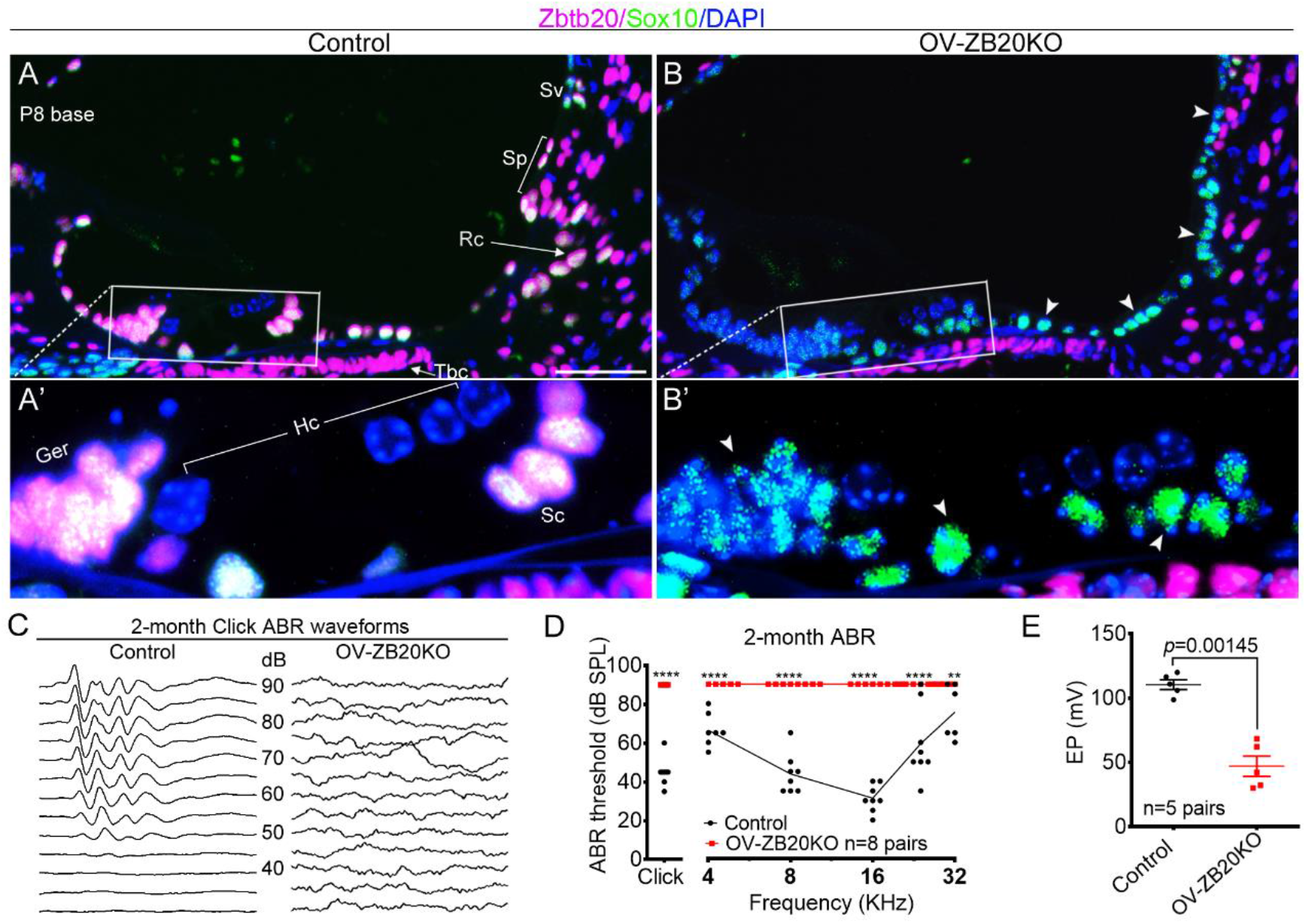
Conditional knockout of *Zbtb20* causes endolymph potential reduction and deafness. (**A and B**) Immunofluorescence staining confirming ZBTB20 protein (magenta) was detected in Sox10-positive (green) supporting cells (Sc), greater epithelial ridge (Ger), root cells (Rc) and spiral prominence epithelial cells (Sp) from control (**A**) but not from OV-ZB20KO (**B**) at P8. Arrowheads indicates negative staining in OV-ZB20KO. (**A’**) and (**B**’) are magnified views of indicated boxed regions in (**A**) and (**B**), respectively. Nuclei were stained with DAPI (blue). Scale bars: 50 µm. n=4 mice/group. Hc, hair cell; Sv, stria vascularis; Tbc, tympanic border cell. (**C**) Representative Click-evoked ABR waveforms for control and OV-ZB20KO mice at 2 months (M) of age. Broadband click stimuli were applied at sound pressure levels (SPL) indicated in decibels (dB). (**D**) Averaged thresholds of Click- and Pure tone-evoked ABRs of control and OV-ZB20KO mice at 2 M. A two-way ANOVA followed by post-hoc pairwise tests (Bonferroni method) was used to compare differences in ABR threshold at multiple frequencies between the two groups. **, *p*<0.01; ****, *p*<0.0001. (**E**) Values of endolymph potential (EP) recorded from the basal cochlear turns from control and OV-ZB20KO mice at 2 M. All values are represented as mean ±SEM. EP values were compared by using a paired Student’s t-test (two-tailed). Abbreviations are consistent across figures in this manuscript. Source data are provided in the Source Data File. See also *SI Appendix* Fig. S4 and S5.

Similar to *Zbtb20*^*-/-*^ mice, OV-ZB20KO mice displayed head tilt and circling behavior without startle responses. Click-evoked ABR tests revealed no stimulus-following ABR waveforms for any sound intensity tested in OV-ZB20KO mice at the age of 2 months, whereas age-matched control mice exhibited the characteristic ABR waveforms at sound pressure levels (SPL) as low as 45 dB (Fig. 2*C*). Tone-burst ABR audiograms showed profound hearing loss at all frequencies tested in the mutant mice (Fig. 2*D*). Similar defects were also detected at P16 (*SI Appendix* Fig. S5*A-B*) and P19 (*SI Appendix* Fig. S5*C*), suggesting congenital deafness in OV-ZB20KO mice. To determine the effect of *Zbtb20* deletion on the cochlear electrochemical environment, we further measured EP. Remarkably, 2-month-old OV-ZB20KO mice exhibited a >50% reduction in EP values compared to their WT littermates (Fig. 2*E*).

Collectively, these data suggest that ZBTB20 expression in OV-derived cochlear epithelial cells and SG neurons is required for hearing in mice.

### *Zbtb20* deficiency permanently arrests cochlear maturation

To explore the pathological mechanisms about the deafness in *Zbtb20* mutants, we performed cochlear histological examination. At P0∼P3, there was no significant difference in the expression of molecular markers for HCs and subtypes of SCs between OV-ZB20KO and control cochleae, suggesting that ZBTB20 is not required for early-stage development of HCs and subtypes of SCs (*SI Appendix* Fig. S6).

The first sign of defect was detected in the outer sulcus at P4, when WT cochleae started to show the development of root cells, as evidenced by the presence of several epithelial cells with an elongated spindle-like shape penetrating into the spiral ligament (*SI Appendix* Fig. S7*A*). In contrast, OV-ZB20KO outer sulcus was still covered by a simple layer of cuboidal-shaped epithelial cells with no sign of root cell development (SI Appendix Fig. 7*B*). At P8, OV-ZB20KO cochleae displayed obvious retardation of cochlear maturation and cellular remodeling compared to WT littermates, showing an absence in the formation of the tunnel of Corti and Nuel’s spaces, a persistent cell mass in the GER with a limited opening of the inner sulcus, an apparently thicker TM that was abnormally attached to the underlying GER and LER, initial sign of root cell development in the outer sulcus, and a flattened SP (Fig. 3*A* and *B*). At P10, the tunnel of Corti and the space of Nuel in OV-ZB20KO mice opened partially, suggesting at least a 2-day delay in the remodeling of the organ of Corti compared to control mice (*SI Appendix* Fig. S7*C* and *D*). Surprisingly, a distorted tunnel of Corti was first manifested with an obviously shorter inner pillar cell relative to the outer pillar cell in the same section of the mutants, indicating a lack of developmental synchronization between inner and outer pillar cells (*SI Appendix* Fig. S7*C* and *D*).

**Figure 3.**
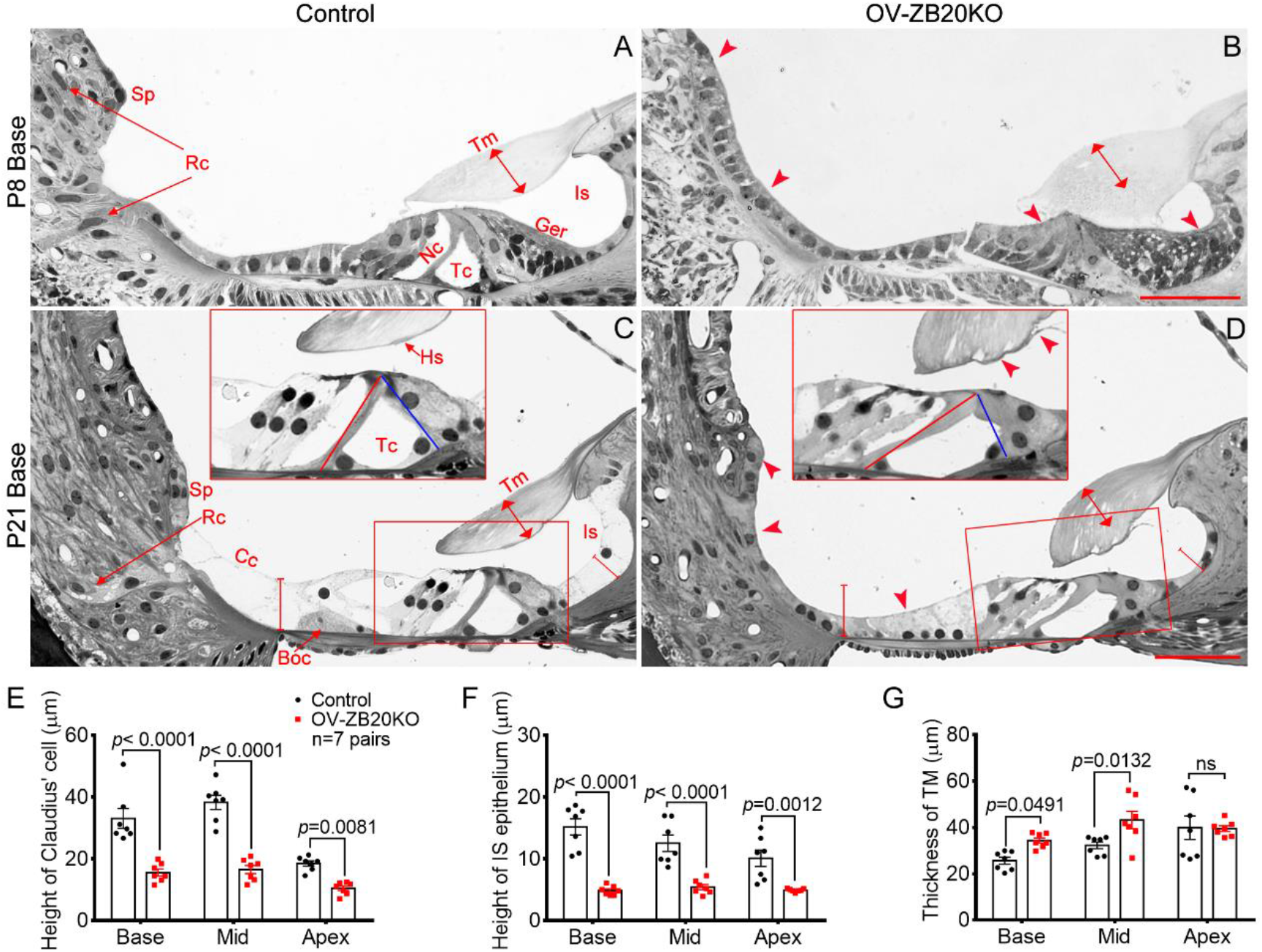
Targeted *Zbtb20* ablation permanently arrests postnatal cochlear development. **(A and B**)Representative semi-thin sections from control and OV-ZB20KO mice at P8, showing the defects (indicated by arrowheads) in OV-ZB20KO cochleae, including the closed spaces of Nuel (Nu) and tunnel of Corti (Tc), and delayed root cell (Rc) development. In addition, the spiral prominence (Sp) appeared flattened, the greater epithelial ridge (Ger) showed initial sign of regression leading to a limited opening of the inner sulcus (Is), and the tectorial membrane (Tm, indicated by the line with arrowheads in both ends) was thickened and abnormally attached to the underlying Corti’s organ. (**C and D**) Representative sections showing the defects in OV-ZB20KO cochleae at P21, including a collapsed tunnel of Corti (Tc) with a deformed inner pillar cell (the red and blue lines indicating the length of the outer and the inner pillar cells, respectively), a thicker and more contracted tectorial membrane (Tm) with deformation at its lower surface around the Hensen’s stripe (Hs), an immature outer border region with no identifiable Boettcher cells (Boc), an immature outer sulcus epithelia with very limited root cells (Rc), and a flattened spiral prominence (Sp). The inserts are magnified images of boxed regions in each image. (**E-G**) Histomorphometry on semi-thin cochlear sections showing significantly decreased height of Claudius cells (Cc) (**E**) and inner sulcus (IS) epithelium (**F**), and increased TM thickness (**G**) in OV-ZB20KO at P21. n=7 mice/group. All values are represented as mean ±SEM. A two-way ANOVA followed by post-hoc pairwise tests (Bonferroni method) was used to compare differences between the two groups. ns, not significant. See also *SI Appendix* Fig. S6-9.

By P21, the organ of Corti displayed a mature structure in control cochleae (Fig. 3*C*). In contrast, OV-ZB20KO cochleae exhibited a severely malformed organ of Corti, similar to that seen around P10, including a misshapen tunnel of Corti, significantly decreased heights in the Claudius’ and inner sulcus cells, and a lack of identifiable Boettcher cells in the basal outer border region (Fig. 3*D-F*). The defects in the mutant outer sulcus and SP were comparable to those seen at P8 (Fig. 3*D*). The TM in OV-ZB20KO cochcleae was significantly thicker in the basal and middle turns, and more contracted in all turns compared with control littermates. In addition, its lower surface around the presumptive Hensen’s stripe, was consistantly delaminated and distorted with an abnormal protrusion (Fig. 3*D* and *G*). The malformed cochlear morphology was largely maintained in adult OV-ZB20KO mice (SI Appendix Fig. S7*E-F*). Thus, deletion of *Zbtb20* throughout the otocyst leads to a permanent disruption in postnatal cochlear development.

Finally, analysis of cochleae from 1-month-old control and OV-ZB20KO mice indicated mild early-onset degeneration of OHCs, which may be a secondary effect of the defect in supporting cells or reflect the cell-autonomous role of ZBTB20 in OHCs, while IHCs remained intact (*SI Appendix* Fig. S8*A*-*C*). No significant differences were detected between control and the mutant group with respect to the number of IHC ribbon synapse identified by RIBEYE/Ctbp2 immunostaining at P10 (*SI Appendix* Fig. S8*D-E*), cochlear innervation at P4 (*SI Appendix* Fig. S9*A-B*), expression patterns of Tuj1(*SI Appendix* Fig. S9*C-D*), Calretinin (a marker for type I neurons) (*SI Appendix* Fig. S9*E-F*), and 3’-cyclic nucleotide 3’-phosphodiesterase (CNPase, a marker for myelinating Schwann cell) in SG at P10 (*SI Appendix* Fig. S9*G-H*), as well as SG neuron density at 2 M (*SI Appendix* Fig. S9*I*-*K*).

Together, these findings suggest that ZBTB20 is required for the terminal differentiation and maturation of cochlear non-sensory epithelial cells.

### *Zbtb20* deletion causes severe defects in root cell differentiation

Given that EP is the main driving force for hair cell mechanotransduction and its reduction causes severe hearing loss or even deafness (30), we next explored the pathological basis for the EP reduction in OV-ZB20KO mice. We first histologically examined the SV, which is required to generate EP and to maintain K^+^ homeostasis in the endolymph (31). No differences in the histology of the SV or adjacent fibrocyte layer were observed between OV-ZB20KO and WT control mice (*SI Appendix* Fig. S10*A-C*). The expression of EP-generating ion-transport molecules and specific markers for three different cell layers were comparable in the SV between the two genotypes (*SI Appendix* Fig. S10*D*), suggesting that the reduced EP in OV-ZB20KO mice is most likely due to the defects beyound SV.

Based on these results, we reasoned that the reduced EP most likely originates from defects in the outer or inner sulcus regions in OV-ZB20KO cochleae. Root cells are located in the outer sulcus region and have been suggested to be involved in K^+^ recycling and establishment of the EP (4, 32-34). To better characterize root cells, OV-ZB20KO and FoxG1-cre mice were crossed to ROSA26-LacZ reporter mice. LacZ staining on cochlear sections showed that root cells in OV-ZB20KO cochlear basal turns had markedly reduced extensions into the spiral ligament by comparison with the extensive long branches observed in control mice (Fig. 4*A-B*).

**Figure 4.**
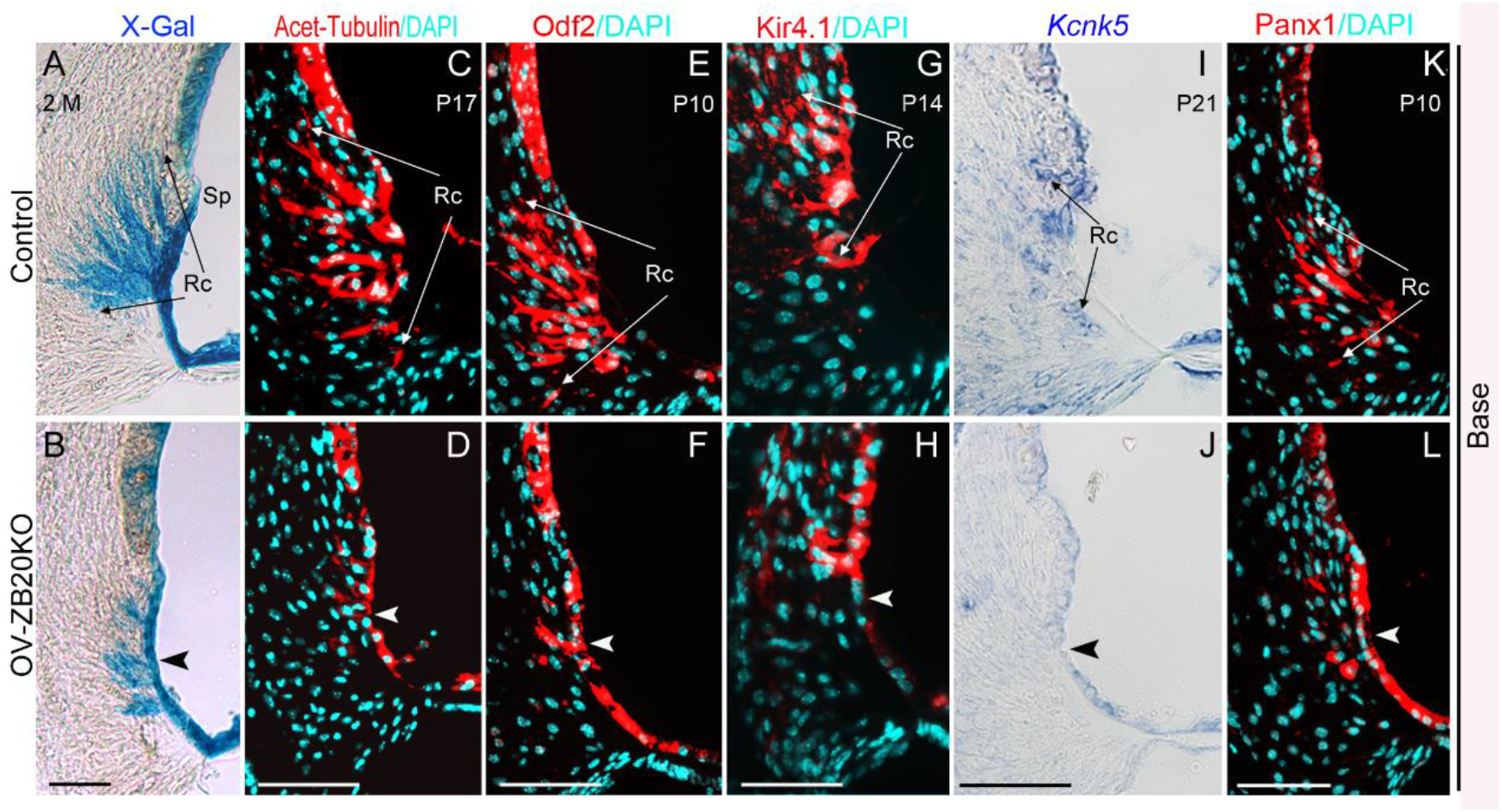
Deletion of *Zbtb20* causes severe defects in root cell development. (**A-B**) X-gal staining (blue) showing root cell processes (Rc) whithin the spiral ligament from control (FoxG1-Cre) and OV-ZB20KO mice crossed with ROSA26-lacZ reporter mice at 2 month (M). The arrowhead indicates very limited root cell (Rc) processes in the mutant mice. (**C-F**) Immunofluorescence staining showing acetylated tubulin (red) and Odf2 (red) expression in the control and mutant lateral wall at indicated postnatal days and cochlear locations. (**G-L**) Kir4.1 protein (red), *Kcnk5* mRNA (blue) and Panx1 protein (red) were detected by either immunofluorescence staining or ISH at indicated postnatal days. Arrowheads in (**D, F, H, J** and **L**) highlight alterations in the expression patterns of indicated molecules in the mutant root cells. Nuclei were stained with DAPI (turqoise). Sp, spiral prominence. Scale bars=50 µm. n=4 animals/group for each marker. See also *SI Appendix* Fig.S10-S11.

The defects of root cell differentiation in OV-ZB20KO were further demonstrated by immunofluorescence staining for acetyl-tubulin and Odf2 (outer dense fiber of sperm tails 2). Consistent with previous observations (35, 36), robust and extensive labeling for acetyl-tubulin and Odf2 were detected in control root cell processes. In contrast, the mutant cochlea displayed the expression of these two molecules largely restricted to the immature outer sulcus epithelium lining the scala media (Fig. 4*C-F*).

We then analyzed expression of ion channels/transporters and gap-junction channel-forming proteins known to be expressed by root cells and closely related to the maintenance of EP and hearing, including KCNJ10 (potassium inwardly-rectifying channel, subfamily J, member 10, also known as Kir4.1)(4), KCNK5 (potassium two pore domain channel subfamily K member 5) (32), connexin26 (Cx26), Cx30 (37) and Pannexin1 (Panx1) (38). In line with previous reports, positive signals for Kir4.1 protein, *KCNK5* mRNA, and Panx1 protein were observed on root cells bodies with their processes penetrating into the spiral ligament in control cochleae, but in OV-ZB20KO cochleae, expression of these molecules was confined to the immature outer sulcus epithelium bordering the endolymph (Fig. 4*G-L*). In contrast, the expression patterns of Cx26 and Cx30 were similar in both control and KO cochleae (*SI Appendix* Fig. S11). Taken together, these results suggest that *Zbtb20* is essential for root cell differentiation. Considering the postulated role of root cells in maintenance of EP, these defects could account for the lowered EP in OV-ZB20KO mice.

### *Zbtb20* deletion disrupts the morphogenesis of spiral prominence

The outer sulcus root cells are closely related to the SP, a buldging structure situated between the upper border of the outer sulcus and the lower margin of the stria. It has been suggested that SP may also play a role in the maintanence of endolymph homeostasis (39), however little is known about the mechanisms underlying its morphogenesis. The abnormal flattened SP observed in OVZB20KO cochlea as shown in Fig 3*A-D* is very intriguing, suggesting ZBTB20 is also essential for the development of this structure.

To better reveal the role of ZBTB20 in the SP development, we examined temporal morphological changes in OVZB20KO SP in greater detail. At P4, the prospective SP in both control and OVZB20KO cochleae appeared as an immature flattened structure covered with several cuboidal epithelial cells, which could be distinguished from the adjacent outer sulcus epithelial cells/root cells by their more darkly stained cytoplasm (*SI Appendix* Fig. S12*A-B*). At P8, coincided with the development of root cells, a buldging SP began to be detected in WT controls, with elongated epithelial cells covering several superiormost root cells. In contrast, OVZB20KO counterparts showed an immature morphology similar to that seen at P4, suggesting a delay in the differentiation of SP epithelium. We then examined the expresion of Anxa1(Annexin A1), a marker for postnatal SP epithelial cells and SV spindle cells (40). At P4 (*SI Appendix* Fig. S12*C-D*) or P10 (Fig. 5*C-D*), strong signals for *Anxa1* mRNA were detected in WT control but not in OVZB20KO SP epithelium, confirming ZBTB20 deficiency disrupting the differentiation of SP epithelium. This was further demonstrated by triple immunofluorescence staining for Anxa1, Epyc (Epiphycan, a marker for root cell) and Wnk4 (a maker for SV basal cells) (Fig. 5*E-F*).

**Figure 5.**
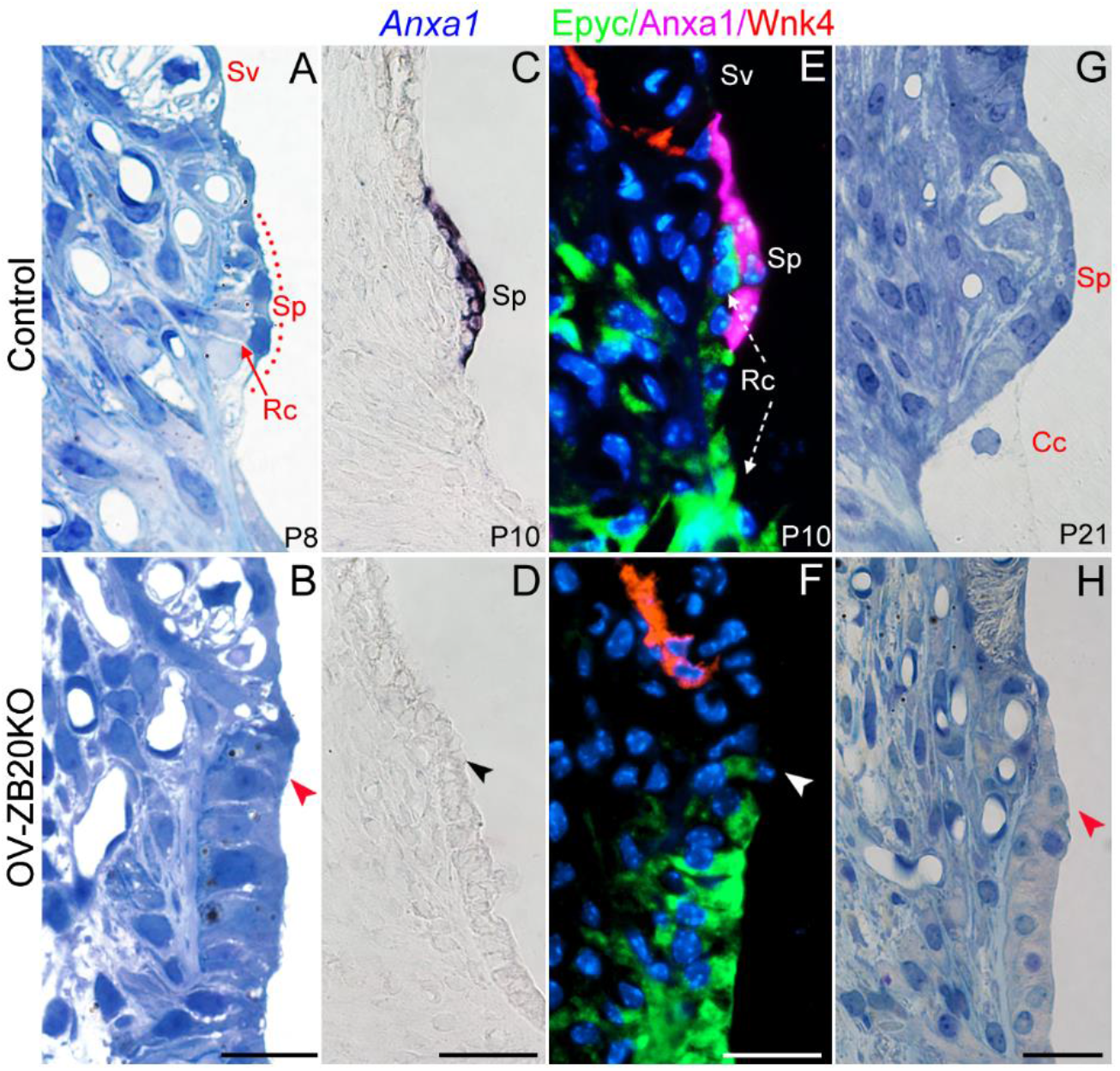
Ablation of *Zbtb20* disrupts postnatal development of spiral prominence. **(A-B)** Representative semi-thin toluidine blue-stained cochlear sections at P8 showing a delay in the differentiation of spiral prominence (Sp) epithelial cells and root cells (Rc) in OV-ZB20KO mice compared with those in control mice at P8. Sv: Stria vascularis. **(C-D)** In situ hybridization showing Anxa1 mRNA was detected in WT control but not OV-ZB20KO spiral prominence epithelium at P10. **(E-F)** Representative immuno-stained cochlear sections showing an absence of Anxa1 expression (magenta) in OV-ZB20KO spiral prominence epithelium at P10. SV was demarcated by Wnk4-positive (red) basal cells. Root cells were stained with anti-Epyc (green). Nuclei were stained with DAPI (Blue). Of note, Anxa1-positive Sp epithelial cells covered the underlying Epyc-positive root cells (Rc) in WT control. **(G-H)** Representative semithin cochlear sections showing a flattened spiral prominence in OV-ZB20KO mice at P21 while WT counterparts exhibiting a mature morphology with squamoid epithelial cells directly connected with Claudius cells (Cc, indicated by black dotted line) covering the underlying root cells. Scale bars: 20 µm in (**A**-**B** and **E**-**H**), 50 µm in(**C**-**D**). n=4 mice/group. See also *SI Appendix* Fig.S12.

At P21, the buldging SP in WT controls exibited a mature morphological feature, showing a squamoid epithelial layer directly connected with the superiormost Claudius cells covering the underlying root cell processes. In contrast, OVZB20KO counterpart remained flattened, with an immature appearance in both SP and the adjenct outer sulcus epithelium similar to that seen at P4 (Fig. 5*G-H*). Accordingly, the expression of *Anxa1* was absent in the mutant SP epithelium (*SI Appendix* Fig. S12*E-F*). These defects were largely maintained at 2 months (*SI Appendix* Fig. S12*G-J*), supporting a failure in the maturation of SP epithelium in the mutant.

### *Zbtb20* deletion does not affect the lineage commitment of cochlear lateral epithelium

The above findings reveal that *Zbtb20* deletion causes permanently arrested development of cochlear lateral epithelium consisting of Claudius, Boettcher, root cells and SP epithelial cells. To rule out the posibility that these defects is due to a lineage commitment failure or cell fate transformation, we examined expression patterns of a set of molecular markers for cochlear lateral region. *Fgfr2* (fibroblast growth factor receptor 2) mRNA is expressed in cochlear lateral non-sensory regions since E13.5 and confined to the outer sulcus around postnatal stage (41). *Fgf16* mRNA is expressed very specifically in the area of the developing SP at embryonic stage (42), while Bmp4 (Bone morphogenetic protein 4) is expressed in the developing lateral border region at postnatal stage (43). There was no significant difference between mutants and controls with respect to the expression patterns of *Fgfr2, Fgf16* at E15.5 and *Bmp4* at P4 (Fig. 6*A-F*). Furthermore, OVZB20KO showed expression domains of *Slc26a4, Lgr5* and *Fgfr2* mRNA (markers for outer sulcus epithelial cells/root cells)(44) confined in the outer sulcus epithelium lining the scala media at P4 (Fig. 6*G-L*), though lacking an extension of those domains within the spiral ligament (corresponding to the developing root cells) observed in WT controls. Together, these data support that the terminal differentiation of cochlear lateral epithelial cells were disrupted but their cell lineages were unaffected by the absence of ZBTB20.

**Figure 6.**
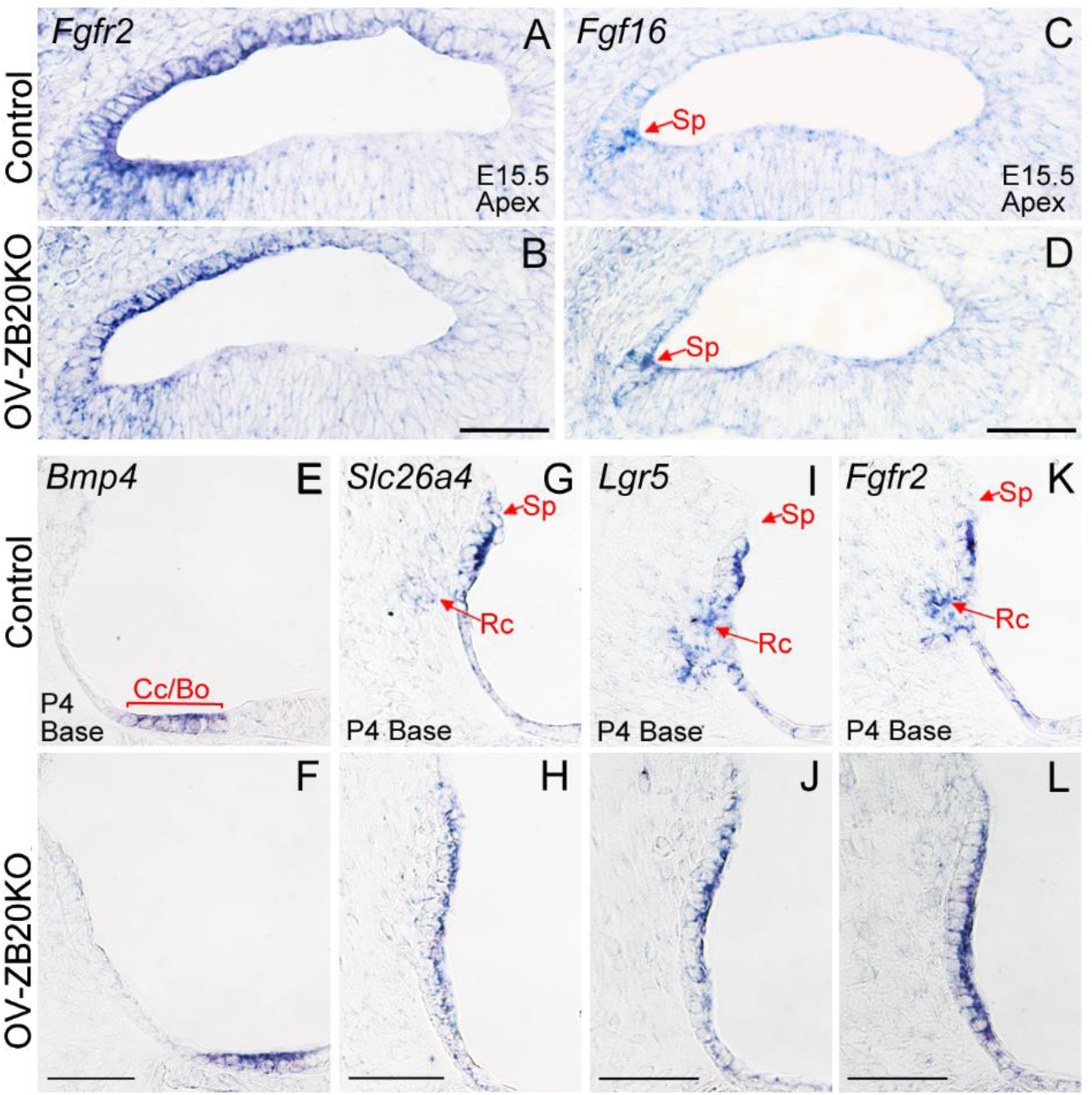
*Zbtb20* deletion does not affect the lineage commitment of cochlear lateral epithelium. **(A-F)** In situ hybridization showing comparable expression patterns of *Fgfr2, Fgf16* and *Bmp4* mRNA in OV-ZB20KO and their control cochleae at indicated ages. Sp, spiral prominence; Cc, Claudius cells; Bo, Bottcher cells. (**G-L**) In situ hybridization showing expression domains of *Slc26a4, Lgr5* and *Fgfr2* mRNA were confined in the outer sulcus epithelium lining the scala media in OV-ZB20KO at P4, though lacking an extension of those domains within the spiral ligament (corresponding to the developing root cells, Rc) observed in WT controls. Scale bars: 50 µm. n=4 mice/group.

### ZBTB20 programs gene expression for cochlear maturation

To characterize the molecular mechanisms underlying the effects of deletion of *Zbtb20* and to elucidate potential downstream targets of ZBTB20, we performed RNA-seq analysis (five biological replicates for each group) on cochlear tissue obtained from OV-ZB20KO and their littermate WT controls at P10, when defects are evident in the mutant. Comparisons between the mutant vs the WT groups identified 155 differentially expressed genes (DEGs) based on a fold change of ≧2 and Q-value≤0.05 (Genes of FPKM<5 were filtered out, with the exception of *Kcnk5*). Seventy-four genes are significantly up-regulated while 81 genes are significantly down-regulated (Fig. 7*A*).

**Figure 7.**
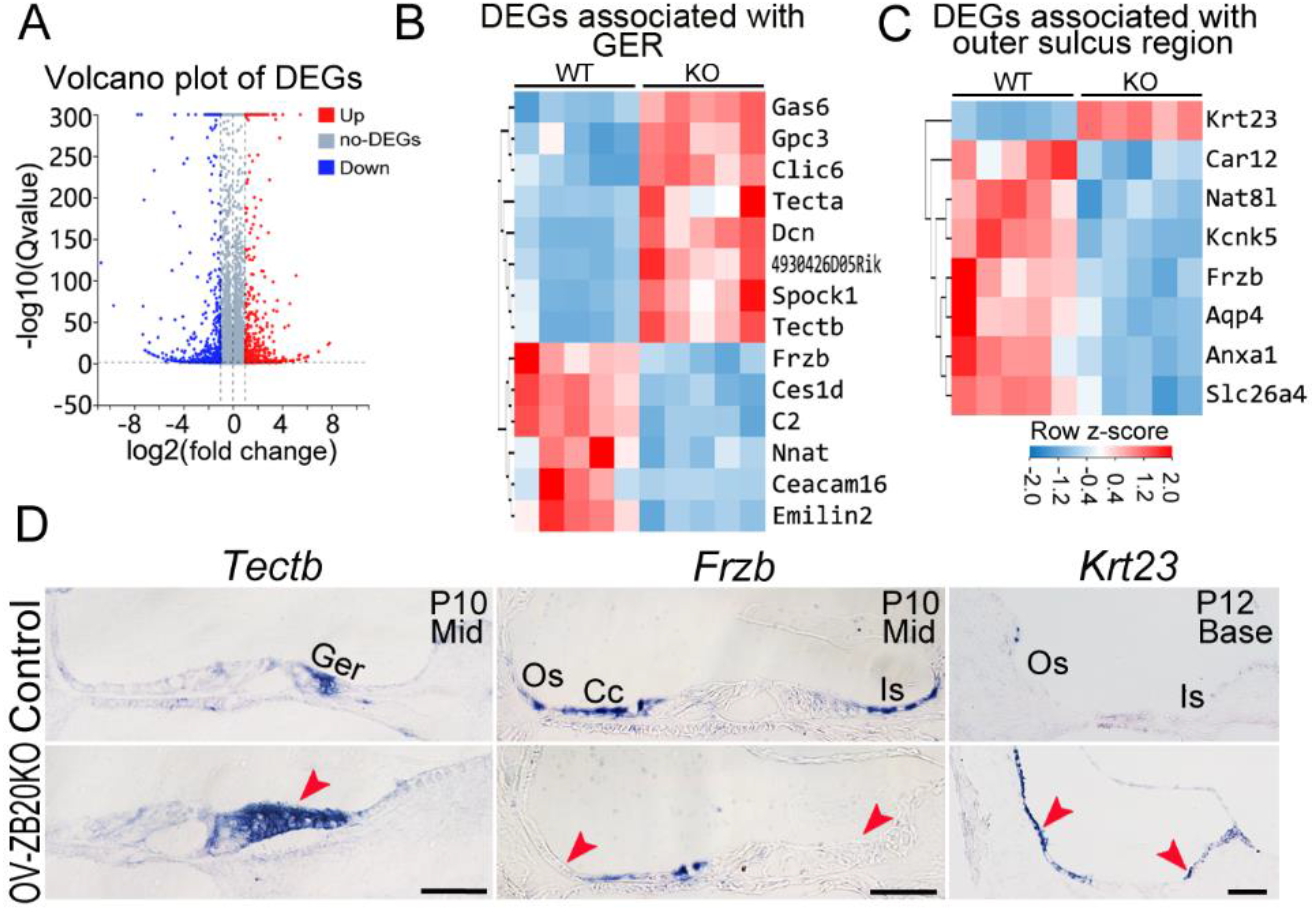
ZBTB20 regulates gene expression for cochlear maturation. (**a**) Volcano plot showing the significantly differentially expressed genes (DEGs) detected in cochlear tissues from P10 OV-ZB20KO compared to WT controls by RNA-seq analysis (n=5 pairs, fold change ≧2, Q value≦0.05). (**B and C**) Heatmap of the DEGs associated with GER (B) or outer sulcus region (**C**) in OV-ZB20KO Vs WT control. (**D**) Validation of DEGs by in situ hybridization analysis. Scale bars: 50 µm. n=3 mice/group. Ger, greater epithelial ridge; Os, outer sulcus; Cc, Claudius cells; Is, inner sulcus.

Fourteen DEGs (8 up-regulated and 6 down-regulated) were known to be predominantly expressed by the GER(45) (Fig. 7*B*). The 8 up-regulated genes mainly code for extracellular proteins including *Tecta* (α-Tectorin) and *Tectb* (β-tectorin), which are the major non-collagenous proteins of the TM (46). The 6 down-regulated genes have been shown to be preferentially expressed by GER at P7 or later but not at P1 including *Frzb* (frizzled related protein, also known as Sfrp3), *Ceacam16* and *Emilin2* (with the exception of *Nnat* (*Neuronatin*), which is expressed by inner sulcus epithelium) (45, 47). The alterations in the gene expression in the GER are consistent with a delay in the remodelling of GER and the increased TM thickness in OV-ZB20KO cochleae.

We also identified 7 down-regulated DEGs associated with outer sulcus region including *Aqp4* (Aquaporin 4), *Kcnk5, Slc26a4*, and *Car12* (Carbonic anhydrase 12), which are predominantly expressed by the outer sulcus/root cells(5, 32, 48-50),(51), and Anxa1, a specific marker for SP epithelium (Fig.7*C*). Unexpectedly, we identified up-regulation of *Krt23* (Keratin 23) in the mutant inner and outer sulci/root cells, which has not been reported to be expressed in cochlea(Fig.7*C-D*). These results are consistent with defects in the differentiation of outer sulcus/root cells and SP epithelial cells.

Among DEGs, 18 were selected for further validation by qRT-PCR (SI Appendix Fig. S15a-d) or in situ hybridization (Fig. 7*E*). These genes were chosen based on either their high fold change or their potential roles in cochlear function.

## Discussion

We and others have previously reported that Zbtb20 functions primarily as a transcriptional factor that mediates developmental processes in a number of different organ systems (16, 17, 21, 52-54). Our findings in this study uncovers an important role of ZBTB20 in cochlear maturation and hearing.

A particularly striking phenotype in OV-ZB20KO cochleae was a significant disruption in the development of root cells in the lateral wall. While the physiological roles of root cells are not well-understood, it is generally accepted that their primary roles are for maintenance of endolymph homeostasis and generation of EP (5, 32). Consistent with this hypothesis, EP first appears around P5 in mice (55), which parallels the development of the root cells. The root cells are located at the lateral border of an epithelial gap junction network within lateral cochlear cells (56). Since gap junctions do not span the basement membrane between the root cell processes and the enveloping type II fibrocytes and/or capillaries, it has been suggested that root cells may release K^+^ into the extracellular space, which is taken by type II fibrocytes, and then relayed back to the SV via the connective tissue gap junction network (57). This hypothesis is supported by the existence of weakly rectifying K^+^ currents through the basolateral processes of root cells (4), the polarized expression of key ion channels/transporters in root cell processes including KCNJ10 (Kir4.1) (58) and KCNK5 (32), both of which are indispensable for the maintenance of EP and hearing. Therefore, the data presented here are consistent with a role for ZBTB20 in the development and function of root cells.

Another intriguing finding of our analysis is the failure in SP morphogenesis upon *Zbtb20* deletion. Although the function of the SP remains largely unknown, it has been proposed to serve as a balance between the perilymph and the endolymph (59). During postnatal development stage, we observed that the SP region has not acquired its bulging appearance until around P8, when immediately below this region an indentation appears (Fig. 5*A*). This indentation is most likely left by the developing root cells which have infiltrated into the spiral ligament and extended their processes towards and beyond SP. Simultaneously, the SP epithelial cells become elongated and cover the superiormost root cells. Based on this observation, we propose that a failure in the morphological transformation of SP epithelial cells together with the defects in root cells development most likely account for a flattened SP upon the deletion of *Zbtb20*. It has to be noted that Claudius cells also undergo dramatic transformation at the same time and extend towards the inferior end of SP. At mature stage, SP epithelial cells form tight junctions with Claudius cells to seperate the underlying root cells from endolymph (57). Thus, the postnatal development of Claudius, root cells and SP epithelial cells must be coordinated to form a normal structure in this region. Our findings support that ZBTB20 plays an essential role in synchronizing this region’s development, the disruption of which leads to defects in fluid homeostasis and EP.

However, it is important to consider that a 50% reduction in EP levels may be not sufficient on its own to cause the level of deafness observed in *Zbtb20*^-/-^ or OV-ZB20KO mice. Instead, deafness in both lines may result from a combination of decreased EP and malformed TM. The TM plays essential roles in cochlear amplification and control of IHC hearing sensitivity (60-62). Normal TM function relies on proper structural formation, in particular on the lower surface, which contains both Kimura’s membrane and Hensen’s stripe, the latter has been reported to either directly contact or to be closely related to stereociliary bundles of IHC (63). Therefore, the disruption in Hensen’s stripe in OV-ZB20KO TM may impair IHC sensitivity (60). In addition, the TM was abnormally thicker and more contracted in OV-ZB20KO cochleae, suggesting that OHCs and their stereocilia might not be properly aligned with Kimura’s membrane, causing an impairment in the amplification of auditory stimuli.

TM defects following deletion of *Zbtb20* most likely originate from a delay in the regression of the GER and maturation of SCs, as the TM is composed of several types of collagen and noncollagenous glycoproteins including Tectα and Tectβ, which are produced by cells in the GER and the developing organ of Corti (46). The delayed GER regression prolongs the expression of tectorins (*Tectα* and *Tectβ*), leading to an increase in the total period of TM growth, likely resulting in a pathological thickening of the TM. Moreover, we noticed that undersurface malformations of the TM were first detected at around P10, when the lower fibrous layer of the TM remained abnormally anchored to the apical surfaces of disproportionately small inner pillar cells. As immature pillar cells also express *Tectα, Tectβ*, and other glycoproteins that are incorporated into the TM (46, 64, 65), it is possible that an arrest of inner pillar cell development may contribute to undersurface deficits in the TM of OV-ZB20KO mice. In addition, the downregulated expression of *Ceacam16* revealed by RNA-seq may also contribute to TM malformation and the disruption in Hensen’s stripe in the mutant, as Ceacam16 can interacts with tectorins and is essential for the formation of Hensen’s stripe (47).

In summary, our data establish an essential role of ZBTB20 in postnatal cochlear development and hearing perception, and particularly the development of cochlear lateral non-sensory epithelium.

## Materials and Methods

### Generation of OV-ZB20KO Mice

*Zbtb20*^-/-^, *Zbtb20*^flox^ mice have been previously constructed in our laboratory (16, 17). OV-ZB20KO mice were generated by crossing *Zbtb20*^flox^ mice with FoxG1-Cre mice (29). Mice have been backcrossed to C57BL/6 mice for at least five generations. Additional mutants were produced by interbreeding Cre-positive heterozygous *Zbtb20*^flox^ and Cre-negative homozygous *Zbtb20*^flox^ mice. Age-matched floxed/Cre-negative littermate mice served as wild-type controls. For some experiments, Cre-positive heterozygous *Zbtb20*^flox^ mice were also crossed to ROSA26-LacZ reporter mice(66). Genotyping was described previously (17). The day when the vaginal plug appeared in the dam was designated as embryonic day (E) 0.5 (E0.5) while the morning following birth was considered as postnatal day 0.5 (P0.5). Data were collected from a single ear per animal for all related experiments. Animal experiments were conducted in accordance with the guidelines of the Naval Medical University Animal Ethics Committee.

### Auditory brainstem response (ABR) test

ABR tests were performed according to a previously reported method (67) with a detailed protocol described in *SI Appendix* Extended Methods.

### Endocochlear potential recording

EP was recorded according to a previously described method (8) with a detailed protocol described in *SI Appendix* Extended Methods.

### Immunohistochemistry and X-Gal staining

Immunostaining was conducted on formalin-fixed, decalcified, paraffin-embedded cochlear mid-modiolar sections at indicated developmental stages according to previously reported methods (18, 21). For cochlear whole-mounts, staining was performed using standard methods as previously described (68). *SI Appendix* Table S1 shows the information for all antibodies used. The detailed protocol for triple immunofluorescence staining was described in *SI Appendix* Extended Methods. F-actin was detected by labeling with Alexa 488-conjugated phalloidin (Invitrogen, 1:100). Phalloidin labelled whole-mount cochleae from 7 mice per genotype were micro-dissected and used to quantify cochlear hair cell loss. All fluorescence images were re-colored using a color blind-friendly palette. X-gal staining was performed as previously described (17). Cochleae from n≥4 mice per genotype at a given age were analyzed.

### In situ hybridization

Mice were first transcardially perfused with 4% paraformaldehyde (pH 9.0). Either embryonic heads or dissected cochleae were then post-fixed, decalcified, and cryostat sectioned at a thickness of 16 μm. Sections were then processed for in situ hybridization as described previously (18). *SI Appendix* Table S2 shows the sequence information for all cRNA probes. Specificities of antisense cRNA probes have been tested. Cochleae from n≥4 mice per genotype at a given age were analyzed.

### Histology and electron microscopy

Mice were first transcardially perfused with 4% paraformaldehyde (pH 7.4). Cochleae were then dissected and gently perfused with 2.5% glutaraldehyde and immersed in the same fixative overnight, followed by post-fixed with 1% osmium tetroxide (OsO4). After decalcification and dehydration, cochleae were embedded in Spurr’s low-viscosity resin and semi-thin (1 µm) mid-modiolar sections were cut and stained with 1.25 % (w/v) toluidine blue. Cochleae from n≥5 mice (n=6∼8 mice if a quantitative analysis was applied) per genotype at a given age were analyzed. The detailed protocol for histomorphometry was described in *SI Appendix* Extended Methods. For SEM analysis, cochleae were prepared as previously described (69).

### Quantitative analysis of ribbon synapses under inner hair cells

Cochlear whole mounts from Control and OV-ZB20KO mice at P10 were immunostained with anti-CtBP2 antibody (BD Biosciences, 612044). Images were obtained and CtBP2 spots (ribbons) were measured with a protocol described in *SI Appendix* Extended Methods.

### RNA-seq

Total RNA was extracted from bilateral cochleae of each individual mouse at P10 (WT control Vs OVZB20KO, 5 biological replicates for each group) using Trizol combined with Qiagen RNeasy Micro kit. The detailed protocol for RNA-seq was described in *SI Appendix* Extended Methods. Significant Differentially Expressed Genes (DEGs) between WT control and OV-ZB20KO cochleae were analyzed using DESeq2 (v1.4.5) based on |log2FC|≥1 and Q-value≤0.05, and genes with FPKM<5 in both groups were filtered.

### Real-time RT-PCR

Total RNA was extracted from bilateral cochleae of each individual mouse (n≥7 mice per genotype). Both on-column and off-column Dnase 1 digestion were applied to eliminate any contaminating genomic DNA. cDNA was synthesized and amplified in duplicate using the SYBR Green PCR assay. RNA expression levels were normalized to those of internal control 36B4 (Rplp0) and presented as fold changes compared with those of the WT Control. The primers used are listed in *SI Appendix* Table S3.

### Statistical analysis

All statistical analysis was performed using GraphPad Prism 7 software. Values are shown as means ±SEMs. Comparisons of two groups (WT Vs OV-ZB20kO) were analyzed using Student’s t test (two-tailed) or a two-way ANOVA followed by post-hoc pairwise tests (Bonferroni method) after the data having passed a normality test. *p*≤0.05 was considered statistically significant. The nonsignificant difference between samples is indicated as “ ns” ; *p* values of significant differences are shown in the graphs.

## Data availability

RNA-seq data are accessible under the accession number of GSE196199. Any remaining data that support the results of this study are available from the corresponding author upon reasonable request.

## Acknowledgements

We thank Sun LH (Xinhua Hospital) for her technical help in ABR tests.This work was supported by grants from the National Key R&D Program (2019YFA0802500, 2018YFA0800602), National Natural Science Foundation of China (31730042, 81771021, 32271162, 91849110, 81472590), and Collaborative Innovation Program of Shanghai Municipal Health Commission (2020CXJQ01).

