## Supporting information for "ZBTB20 is Essential for Cochlear Maturation and Hearing in Mice"

**This PDF file includes:**

Supporting text
Tables S1 to S3
Figures S1 to S13

### Supporting Text

### Extended Methods

#### Auditory brainstem response (ABR) test

Age- and sex-matched mice were anesthetized using an intraperitoneal injection of pentobarbital sodium (50 mg/kg) and kept on a heated mat inside a sound-attenuated chamber. Electrodes were placed subcutaneously over the vertex of the head (active), the left mastoid (reference), and the right hind limb (ground). Click stimuli or pure tone stimuli from 4 to 32 kHz were generated by Tucker Davis Technologies System (TDT) and delivered by a MF1 speaker (TDT). Auditory thresholds were determined as the lowest sound intensity that produced a reproducible ABR response. Data were collected from a single ear per animal. Since no obvious bias in the hearing phenotype between male and female groups were noted (data not shown), the data collected from both sexes were combined and analyzed.

#### Endocochlear potential recording

Tracheotomy was performed on mice in the ventral position after anesthesia. The tympanic bulla was opened and a small hole was made in the cochlear lateral wall in the basal turn using a fine drill. The microelectrode filled with 150 mM KCl was inserted into the scala media through the drilled hole by controlling movement of the micromanipulator. The ground electrode was placed in the dorsal neck muscles. The response from the microelectrode was amplified using an Axopatch 200B amplifier (Molecular Probe, Sunnyvale, CA, USA) and acquired by software pClamp 10

(Molecular Devices). The voltage changes during a penetration were continuously recorded. Single ear was analyzed for each animal.

**Immunohistochemistry**

The primary antibodies used were listed in Table S1. For single immunostaining, signals were amplified and visualized using TSA-Alexa Fluor 594 or 488 or the TSA system combining with streptavidin-horseradish peroxidase (HRP) (BD Company). For double immunofluorescence staining, the second primary antibody was detected using either Alexa 594 or 488-conjugated secondary antibodies (Invitrogen, 1:500). For triple immunofluorescence staining, anti-Epyc (Atagenix, PA8263H) was visualized using TSA-FITC followed by boiling with a striping buffer (PMID: **29739626**), anti-Wnk4 (Proteintech, 22326-1-AP) and anti-Anxa1 (Proteintech, 66344-1-IG) were then applied and visualized using Alexa 594-conjugated secondary antibody and TSA-Cy5, respectively. Nuclei were labeled with DAPI.

**Histomorphometry**

Semi-thin (1 μm) mid-modiolar sections were cut and stained with 1.25 % (w/v) toluidine blue. Cochleae from n≥5 mice (n=6~8 mice if a quantitative analysis was applied) per genotype at a given age were analyzed. For quantitative analysis, the upper basal, middle and apical turns from three non-consecutive mid-modiolar sections for each cochlea (single ear per animal) were photographed at a 200x magnification and the thickness of TM, heights of Claudius and inner sulcus cells, the thickness and area of SV were measured at defined regions using Image-Pro Plus 6.0 program. Similar protocol was applied for the quantification of the area of Rosenthal's canal, except that SG regions were photographed at a 400x magnification. SG neurons with a visible nucleus were counted within Rosenthal’s canal at three cochlear turns from three sections for each cochlea and densities of SG neurons were calculated by the number of SG neurons divided by the area of Rosenthal’s canal.

**Quantitative analysis of ribbon synapses under inner hair cells.**

Cochlear whole mounts from Control and OV-ZB20KO mice at P10 were immunostained with anti-CtBP2 antibody (BD Biosciences, 612044). Images were obtained with a Lionheart FX automated microscope (Agilent BioTek) using a 100X oil-immersion lens and acquired at 0.4 μm step size in the Z-axis in non-overlapping regions. Maximum intensity projections of Z stacks were converted into 2-D images and CtBP2 spots (ribbons) were measured manually from at least 10 IHCs in the apex, or upper-base (8 animals per group).

**RNA-seq**

RNA-seq libraries were prepared and single end 50 bases reads were generated on DNBSEQ platform (BGI-Shenzhen, China). The clean reads were aligned to the Mus musculus reference genome (GCF_000001635.26_GRCm38.p6) using HISAT2 (v2.0.4) and aligned to reference coding gene set using Bowtie2 (v2.2.5)， then expression levels of genes were calculated by RSEM (v1.2.12).

**Table S1. The information for primary antibodies**

| Antibody | Resource | Dilution |
|---|---|---|
| anti-ZBTB20 monoclonal antibody clone E-11 | Santa Cruz, SC-515370 | 1:2000 |
| anti-Myosin VIIa/Myo7A | Abcam, ab3481 | 1:200 |
| anti-Calretinin | Chemicon, AB5054 | 1:200 |
| anti-alpha 1 Sodium Potassium ATPase | Abcam, ab7671 | 1:2000 |
| anti-p75ntr | Millipore, 07-476 | 1:200 |
| anti-Neurofilament 200 | Sigma, N4142 | 1:200 |
| anti-β-Tubulin (9F3) | Cell Signaling, 2128 | 1:500 |
| anti-CNPase | Thermo Fisher, MS-349-P0 | 1:2000 |
| anti-KCNQ1 | Santa Cruz, SC-10646 | 1:2000 |
| anti-Kir4.1 | Alomone labs, APC-035 | 1:2000 |
| anti-NKCC1 | Invitrogen, PA5-98154 | 1:2000 |
| anti-Acetylated Tubulin | Sigma, T7451 | 1:300 |
| anti-Cenexin1/ODF2 | Abcam, ab43840 | 1:2000 |
| anti-CX26 | Abcam, ab65969 | 1:2000 |
| anti-CX30 | Invitrogen, 33-2500 | 1:2000 |
| anti-Panx1 | Abclonal, A13587 | 1:2000 |
| anti-Sox10 | Abclonal, A15100 | 1:2000 |
| anti-Tuj1 | Novus, MAB1195 | 1:2000 |
| anti-Islet1 | Abclonal, A0871 | 1:2000 |
| anti-Epyc | Atagenix, PA8263H | 1:2000 |
| anti-Wnk4 | Proteintech, 22326-1-AP | 1:500 |
| anti-Anxa1 | Proteintech, 66344-1-IG | 1:2000 |

**Table S2. Sequence information for cRNA probes**

| Gene | Accession # | PCR Primers | Probe length (bp) |
|---|---|---|---|
| *Kcnk5* | NM_021542.4 | F: 5' agggcgcctcttctgtgtcttctac<br>R:5' Agaggccggtttcttcattttcaa | 963 |
| *Spry2* | NM_011897.1 | F: 5' cagatgtgttctaagcctgctg<br>R:5' tcccatagtcaatgacgttctg | 694 |
| *Fgfr3* | NM_001163215.2 | F: 5'ccagcagtggagcttggt<br>R: 5'ccaaagcagccttctcca | 857 |
| *Tectβ* | NM_009348 | F: 5' ctaagcttagcccgtcaactactccttctcct<br>R: 5' gtctagactccccagccatgcagtcttct | 780 |
| *Krt23* | NM_033373.2 | F: 5' Acccgtatactaaggtggcatcagg<br>R: 5' acccattagcgcggagacaaagac | 1038 |
| *Bmp4* | NM_007554.1 266-1172 | F: 5'cctgcagcgatccagtct<br>R: 5'gcccaatctccactccct | 907 |
| *Slc26a4* | NM_011867.4 | F: 5'cagagatgcggcccgagtgttg<br>R: 5'tcccgttcttggttgccctgagtt | 1271 |
| *Frzb* | NM_011356.4 | F: 5'tctctcctgaggccatcg<br>R:5'tgcattctcaatcggggt | 853 |
| *Dct* | NM_010024.3 | F: 5'gggcctgcctttgtcacg<br>R:5'ccaggtcaggccaggtagga | 943 |
| *Lgr5* # | NM_010195.2 | F: 5'cctatttggtagctggctgatcc<br>R: 5'agcaacagagcaatgtgcttcacc | 657 |
| *Anxa1*# | NM_010730.2 | F: 5'gagcccctacccttccttcaatg<br>R: 5' tctgtccccttctcctttctcca | 543 |
| *Fgf16*# | NM_030614 | F: 5'gcggaggtcgggggcgtctttg<br>R: 5'tgggggcccttgttctttgtaatg | 909 |
| *Fgfr2*# | NM_001347638.1 | F: 5'cattggaggctataaggtacg<br>R: 5'tccttgggtttgtctttatcg | 897 |

(# )These cRNA probes were prepared using a PCR-based technique (reference: PMID **25624266)**. The reverse primers include the T7 promoter sequence (GGATCCTAATACGACTCACTATAGGGAG) at the 5' end. The antisense-RNA strand was produced by transcription of the PCR product using T7 RNA polymerase.

**Table S3. Sequence information for qRT-PCR primers**

| Gene | GenBank accession no. | Primers for qRT-PCR |
|---|---|---|
| *Crym* | NM_016669 | F: 5’ccgggggctcacatcaat<br>R:5’tcccgggagtccacatacag |
| *Emilin2* | NM_001357336.1 | F: 5’gaccggaagctggctgacctga<br>R:5’tgggccgaaatcgctgctctt |
| *Gas6* | NM_019521.2 | F: 5’gcatggccaagagcgtgaagtcct<br>R:5’gttggcccgctgctggtgatgc |
| *Nell1* | NM_001037906.2 | F: 5’agggtctgccgaggtcataactt<br>R:5’gtccgctgccacaatcatcat |
| *Ttr* | NM_013697 | F: 5’cagcccatactcctacagcaccac<br>R:5’taggagcaggggagaaaaatgagg |
| *Rplp0 (36B4)* | NM_007475.5 | F: 5’aagcgcgtcctggcattgtctgtg<br>R:5’tggttgctttggcgggattagtcg |
| *Car9* | NM_139305.2 | F:5’cactgttgccctcggacctca<br>R:5’gcgtggctcggaagttcagttgta |
| *Tectα* | NM_001324548.2 | F:5’cgccgcctacgcattcc<br>R:5’agccccgacgcctcctatctt |
| *Tectβ* | NM_009348.4 | F:5’gcatgagaacggcaaagaccaca<br>R:5’acagttcacggggcaggagagc |
| *Spock1* | NM_001166463.1 | F:5’ctgcgcccaaaggtgactgtgct<br>R:5’ttcataccccggtttgtgttctcg |
| *Gpc3* | NM_016697.3 | F:5’cgctttgccgggctacatctgc<br>R:5’ccgttccttgccgccttctgg |
| *Slc26a4* | NM_011867.4 | F:5’cagtcttggcggccgttgtcatt<br>R:5’aagccgaggtccagccccagaa |
| *Ceacam16* | NM_001033419.2 | F:5’ccgggaggtgggcttcgctaat<br>R:5’ctgctcactcacgggccacctgta |
| *Kcnj16* | NM_001252207.1 | F:5’aacccagccacagagaccgtaag<br>R:5’gaaggcaccaaaagaatcaaaatg |
| *Vim* | NM_011701.4. | F:5’caggccaagcaggagtcaaacgag<br>R:5’atcctgcaggcggccaatagtgtc |
| *kcnk5* | NM_021542.4 | F:5’gcacaggcccccaaggatagtta<br>R:5’gaggccggtttcttcattttcaa |
| *Frzb* | NM_011356.4 | F:5’ tcccacccccaccaaatctcctc<br>R:5’ ctgcgccggctgctcttcc |
| *Scin* | NM_001146196.1 | F:5’ tggcggtgactgctacattatcc<br>R:5’ ccggtgcctgcccttctttc |

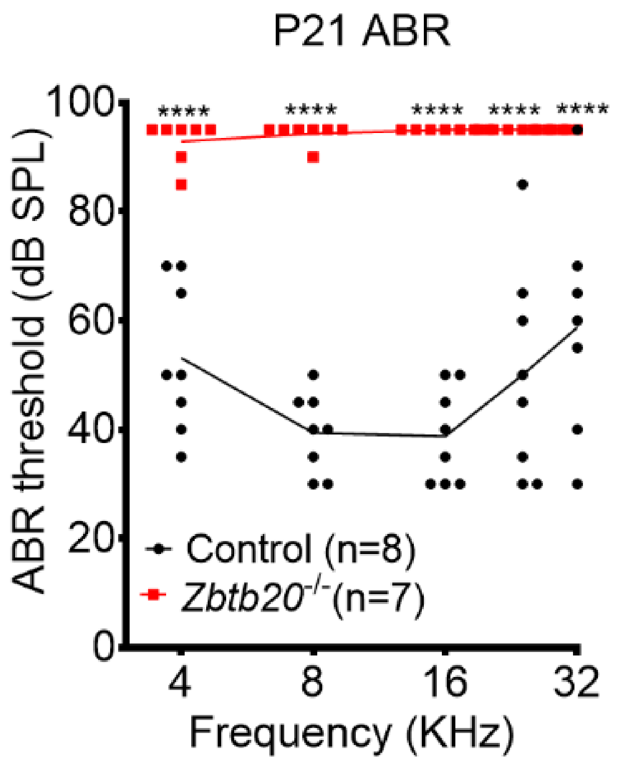


**Fig. S1. Global knockout of *Zbtb20* causes deafness.** Averaged ABR thresholds of WT control and *Zbtb20*$^{-/-}$ mice at P21. Pure-tone stimuli were applied at the frequencies indicated. All values are represented as mean ±SEM. A two-way ANOVA followed by post-hoc tests (Bonferroni method) was used to compare differences between the two groups. ****: P<0.0001.

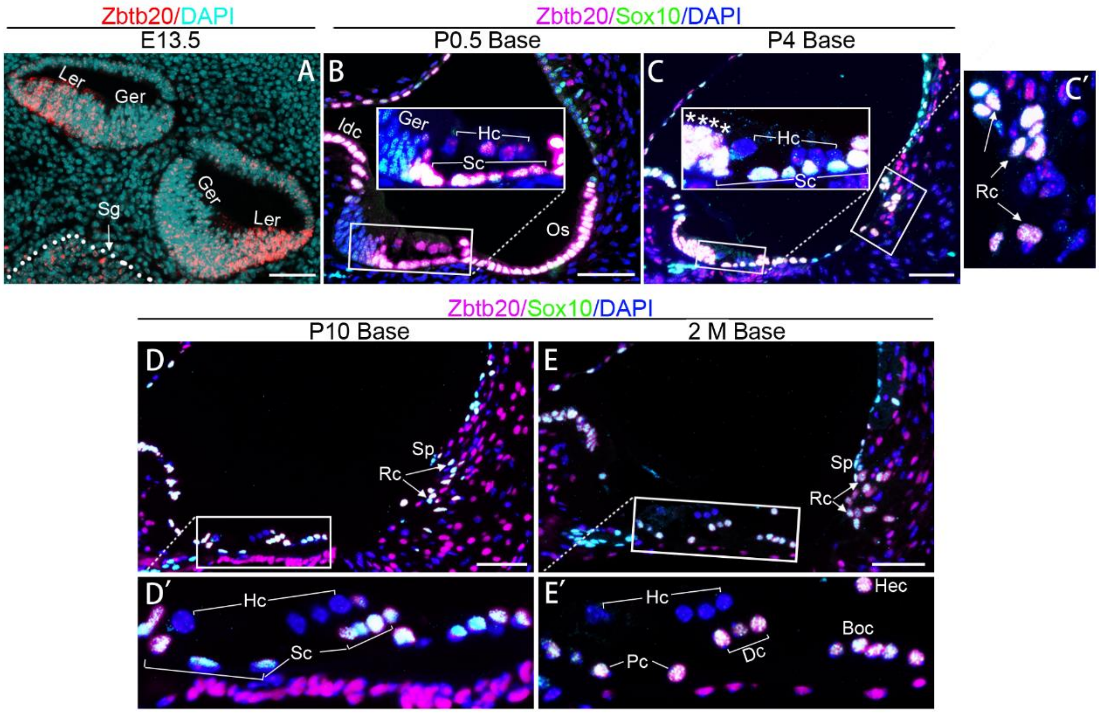


**Fig. S2. ZBTB20 is differentially expressed in cochlear non-sensory epithelial cells.** Immunofluorescence staining showing spatial and temporal expression pattern of ZBTB20 (red or magenta) in cochlea at indicated ages. (**A**) ZBTB20 was detected in cochlear LER and SG region (outlined by white dashed lines) at E13.5. GER showed poor staining at this stage. (**B-E**) ZBTB20 was strongly expressed by Sox10-positive (green) cochlear non-sensory epithelial cells including supporting cells (Sc), interdental cells (Idc), and the outer sulcus (Os)/root cells (Rc) at P0.5 (**B**), P4(**C**), P10 (**D**) or 2 M (**E**). ZBTB20 signals was increased in the lateral and throughout GER at P0.5 or P4, respectively (indicated by *). ZBTB20 signals were not detected in Sox10-negative HCs at P4, P10 or 2 M. (**C', D'** and **E'**) are magnified views of indicated boxed areas in (**C-D**). Inserts are enlarged images of indicated boxed areas in each image. Nuclei were stained with DAPI (turquoise in (**A**), blue in other panels). Pc, pillar cell; Dc, Deiters' cell; Boc, Boettcher cells; Hec, Hensen's cell; Sp, Spiral prominence. Abbreviations are consistent across figures in this manuscript. n=4 mice for each developmental stage. Scale bars in all panels=50 μm.

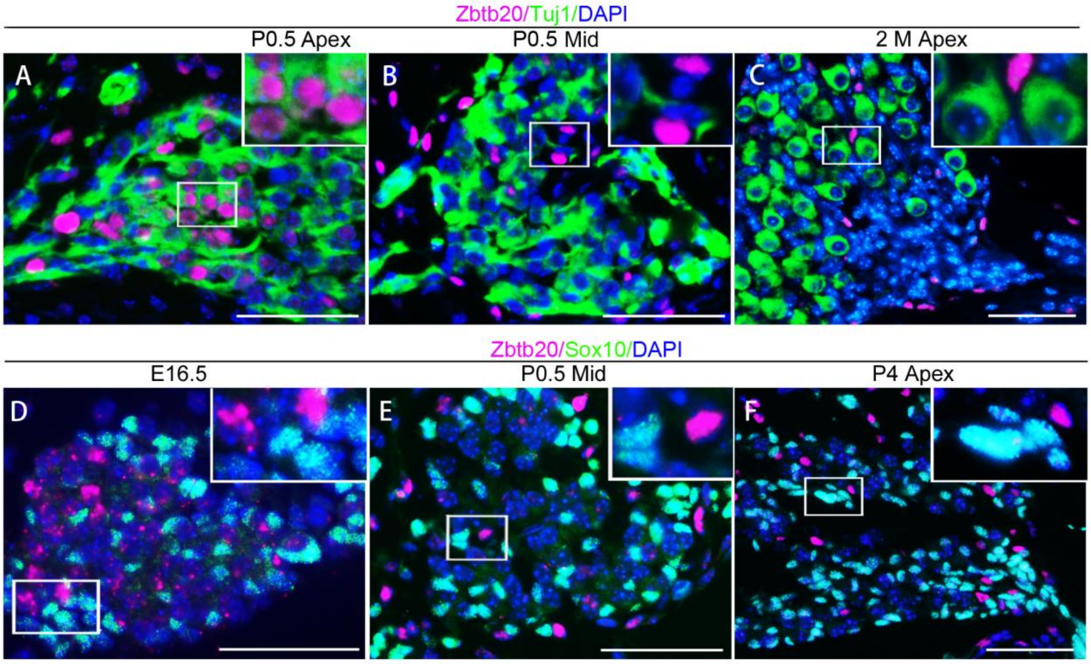


**Fig. S3. ZBTB20 is expressed by a subtype of immature spiral ganglion neurons.** (**A-B**) Double-immunofluorescence staining showing ZBTB20 (magenta) was expressed by a subset of Tuj1-positive (green) spiral ganglion (SG) neurons at apical but not middle turn at P0.5. (**C**) Tuj1-positive SG neurons did not express ZBTB20 at 2 M. (**D-F**) No co-localization of ZBTB20 with Sox10 (green) was observed in SG region at E16.5, P0.5 or P4. Inserts are enlarged images of the boxed area in each image. Nuclei were stained with DAPI (blue). Scale bars in all panels=50 μm.

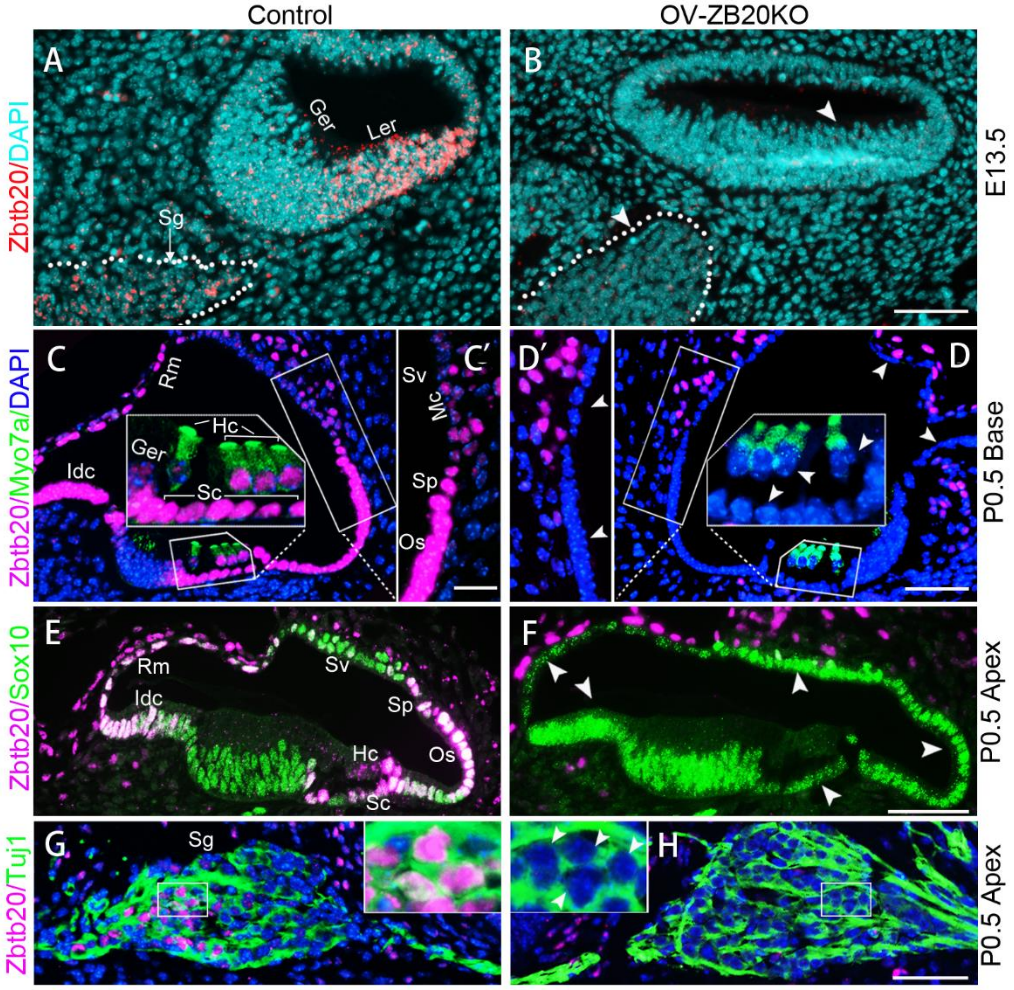


**Fig. S4. *Zbtb20* is specifically deleted in OV-ZB20KO cochlear epithelia and spiral ganglion neurons.** (**A-H**) Immunofluorescence staining showing ZBTB20 (red or magenta) was detected in control (**A, C, E,** and **G**) but not OV-ZB20KO (**B, D, F,** and **H**) cochlear duct and spiral ganglion (Sg, outlined by white dashed lines ) at E13.5 (**A-B**), Myo7a-positive(green) hair cells (Hc) (**C** and **D**), Sox10-positive (green) supporting cells (Sc), interdental cells (Idc); epithelial cells in outer sulcus (Os), spiral prominence (Sp) and Reissner's membrane (Rm); marginal cells (Mc) in stria vascularis (Sv)(**C-F**), as well as a subset of Tuj1-positive (green) spiral ganglion (Sg) neurons (**G** and **H**) at P0.5. Arrowheads in (**B, D, F**, and **H**) indicate negative staining. (**C'**) and (**D'**) are magnified views of indicated boxed regions in (**C**) and (**D**), respectively. Inserts are magnified views of indicated boxed areas in each image. Nuclei were stained with DAPI. Ger, greater epithelial ridge; Ler, lesser epithelial ridge. Scale bars: 50 µm (**B, D, F,** and **G**), 20 µm (**C'** and **D'**). n=4 mice/group.

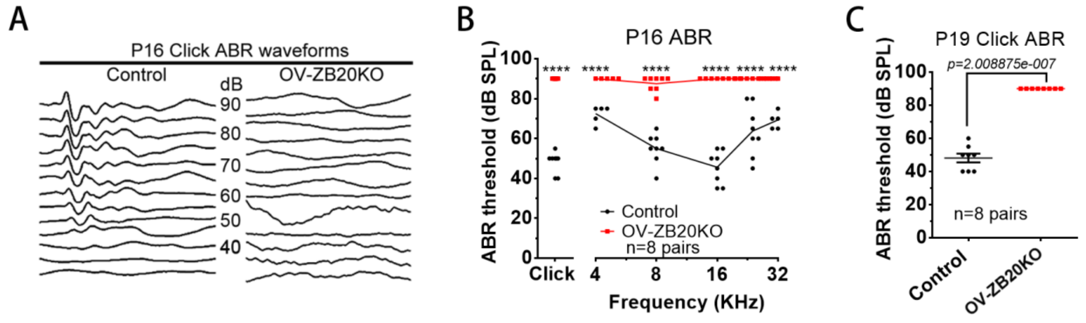


**Fig. S5. Conditional knockout of *Zbtb20* causes deafness in mice.** (**A**) Representative Click-evoked ABR waveforms for control and OV-ZB20KO mice at P16. Broadband click stimuli were applied at sound pressure levels (SPL) indicated in decibels (dB). (**B**) Averaged thresholds of Click- and Pure tone- evoked ABRs of control and OV-ZB20KO mice at P16. (**C**) Averaged thresholds of Click-ABR of control and OV-ZB20KO mice at P19. All values are represented as mean ±SEM. A two-way ANOVA followed by post-hoc pairwise tests (Bonferroni method) was used to compare differences in ABR threshold at multiple frequencies between the two groups. Click-ABR values in (**C**) were compared using a paired Student's t-test (two-tailed). ****: P<0.0001.

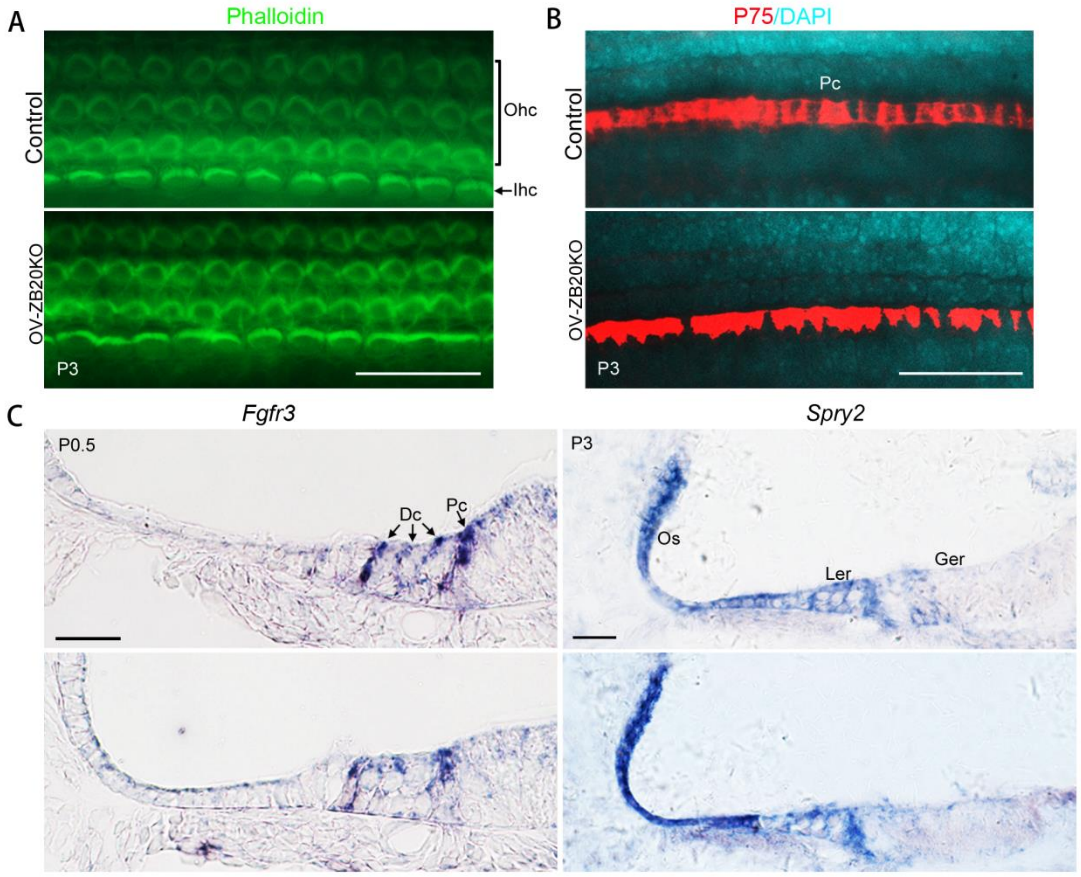


**Fig. S6. Conditional deletion of Zbtb20 does not affect the early-stage differentiation of hair cells or subtypes of supporting cells.** (**A**) Representative cochlear whole mounts stained with phalloidin (green), showing one row of inner (Ihc) and three rows of outer hair cells (Ohc) in both WT control and OV-ZB20KO cochleae at P3. (**B**) Immuno-staining on cochlear whole mounts showing 1 line of p75-positive (red) pillar cells (Pc) in both OV-ZB20KO and WT control at P3. Nuclei were stained with DAPI (turquoise). (**C**) In situ hybridization showing comparable *Fgfr3* and *Spry2* mRNA (blue) expression patterns between WT control and OV-ZB20KO cochleae at P0.5 and P3. Dc, Deiter's cell; Ger; greater epithelial ridge; Ler, lesser epithelial ridge; Os, outer sulcus. Scale bars in all panels=50 μm. n=5 mice/group.

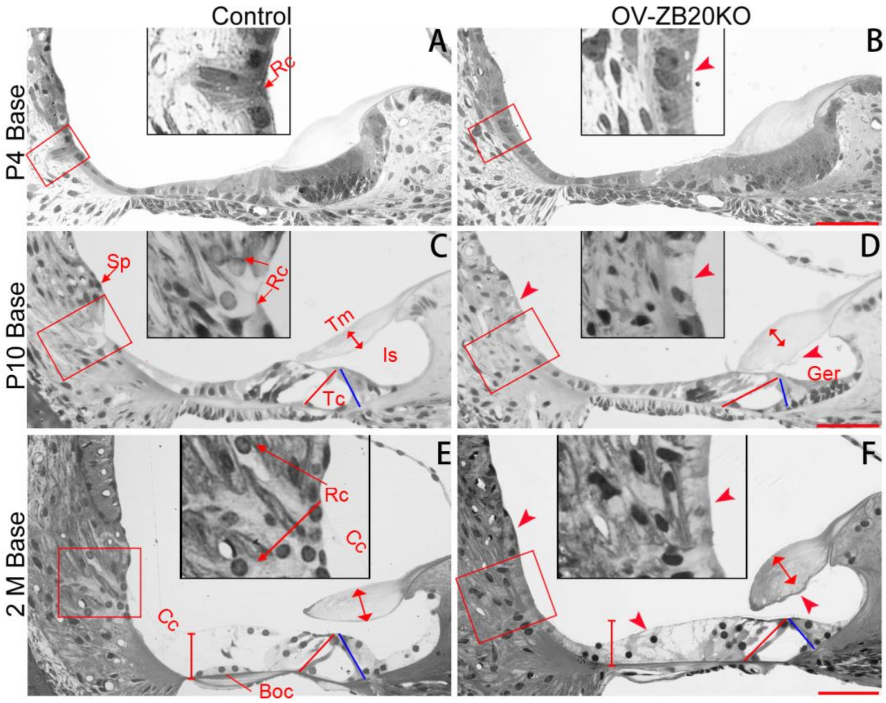


**Fig. S7. Conditional deletion of *Zbtb20* permanently arrests postnatal cochlear development.** Representative semi-thin toluidine blue-stained cochlear sections from the WT control and OV-ZB20KO mice at P4 (**A-B**), P10 (**C-D**) and 2 M (**E-F**). (**A-B**) OV-ZB20KO basal cochleae showing delayed root cells (Rc) development (indicated by the arrowhead) in the outer sulcus region compared to WT control at P4. (**C-D**) OV-ZB20KO cochleae at P10 showing a collapsed tunnel of Corti (Tc) with a deformed inner pillar cell (the red and blue lines indicating the length of the outer and the inner pillar cells, respectively), delayed GER regression with a limited opening of inner sulcus (Is), limited root cells (Rc) development in the outer sulcus. In addition, the spiral prominence (Sp) appeared flattened, and the tectorial membrane (Tm, indicated by the line with arrowheads in both ends) was thickened and abnormally attached to the underlying Corti's organ. (**E-F**) 2-month OV-ZB20KO cochleae showing a collapsed tunnel of Corti (Tc), a thicker and more contracted tectorial membrane (Tm) with deformation at its lower surface, an immature outer border region with no identifiable Boettcher cells (Boc), and very limited root cells (Rc). Inserts are magnified views of boxed areas in each image. Scale bars in all panels=50 μm. n=5 mice per group for each stage.

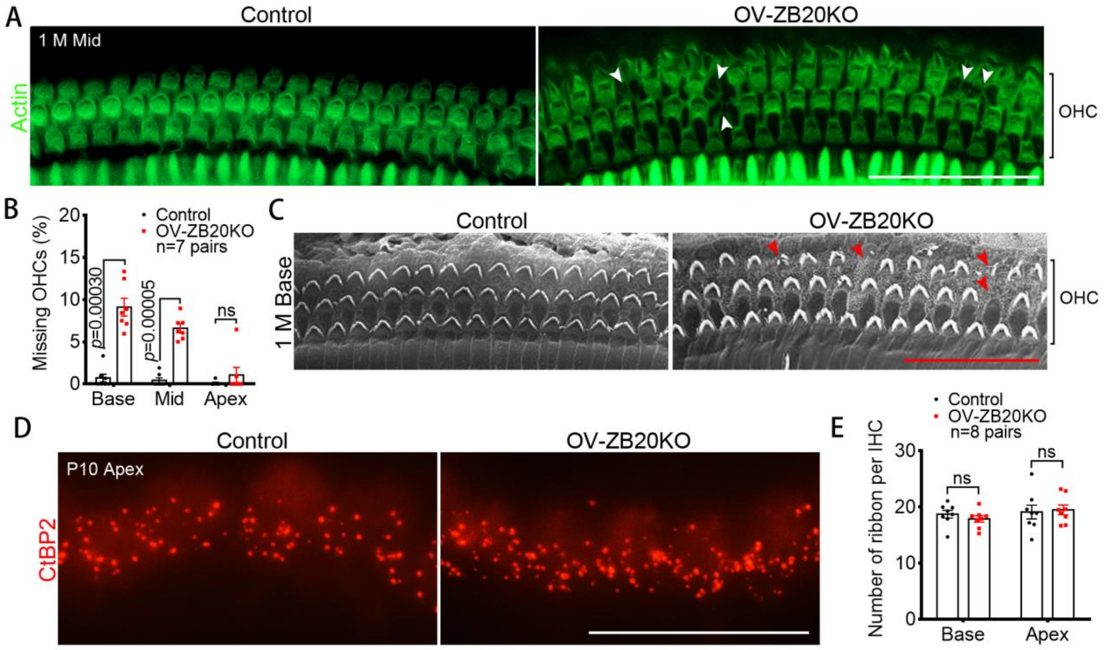


**Fig. S8. Conditional ablation of *Zbtb20* causes early-onset degeneration of outer hair cells.** (**A**) Representative cochlear whole mounts stained with phalloidin (green) showing missing outer hair cells (OHCs) (indicated by white arrowheads) in OV-ZB20KO but not in WT control mice at 1 month (M). (**B**) Quantification of missing OHCs. n=7 mice/group. (**C**) Representative views of 1-month cochleae under the scanning electron microscopy, showing missing OHCs (indicated by red arrowheads) in OV-ZB20KO but not in control mice. n=4 mice/group. (**D**) Representative cochlear whole mounts immuno-stained for CtBP2 (red) showing comparable presynaptic ribbons of inner hair cells between control and OV-ZB20KO mice at P10. (**E**) Quantification of CtBP2-positive ribbons per IHC. n=8 mice/group. Scale bars: 50 μm (**A**), 30 μm (**C**), 20 μm (**D**). Values are represented as mean ±SEM. Paired Student's t-test (two-tailed) was used to compare the values of the two groups. ns: not significant.

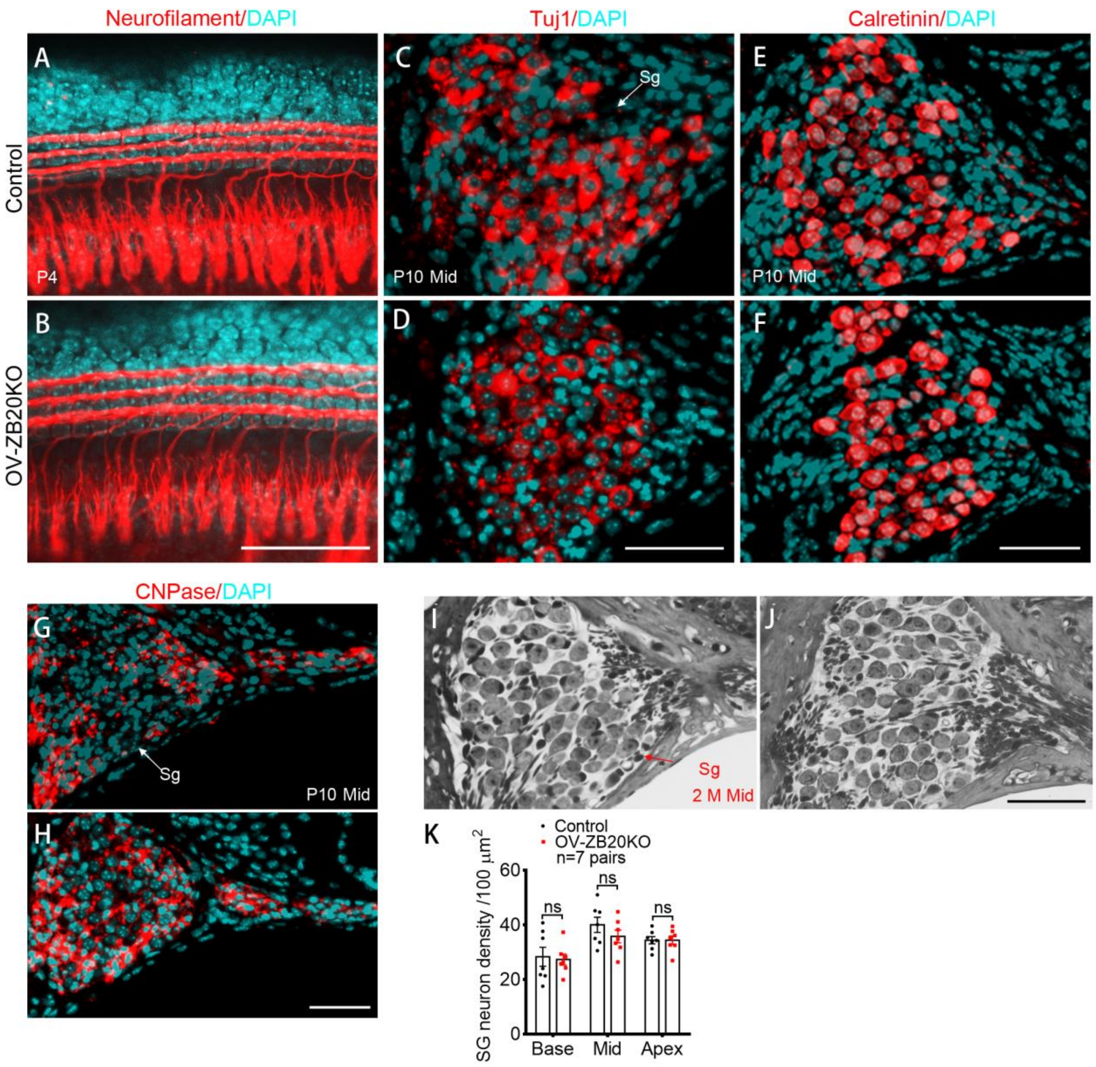


**Fig. S9. Ablation of *Zbtb20* does not affect cochlear innervation and spiral ganglion development.** (**A-H**) Representative views of cochlear whole mounts at P4 (**A-B**) or cochlear SG sections at P10 (**C-H**) immuno-stained with indicated molecular markers (red), showing comparable expression patterns between control and the mutant groups. Nuclei were stained with DAPI (turquois). n=5 animals/group for each marker. (**I-J**) Representative semi-thin toluidine blue-stained cochlear SG sections from control and OV-ZB20KO mice at 2 months (M). (**K**) Graphs showing that densities of SG neurons in OV-ZB20KO cochleae were comparable to those in control at 2 M, n=7 mice/group. Values are represented as mean ±SEM. Paired Student's t-test (two-tailed) was used to compare the values of the two groups. ns: not significant. Scale bars in all panels=50 μm.

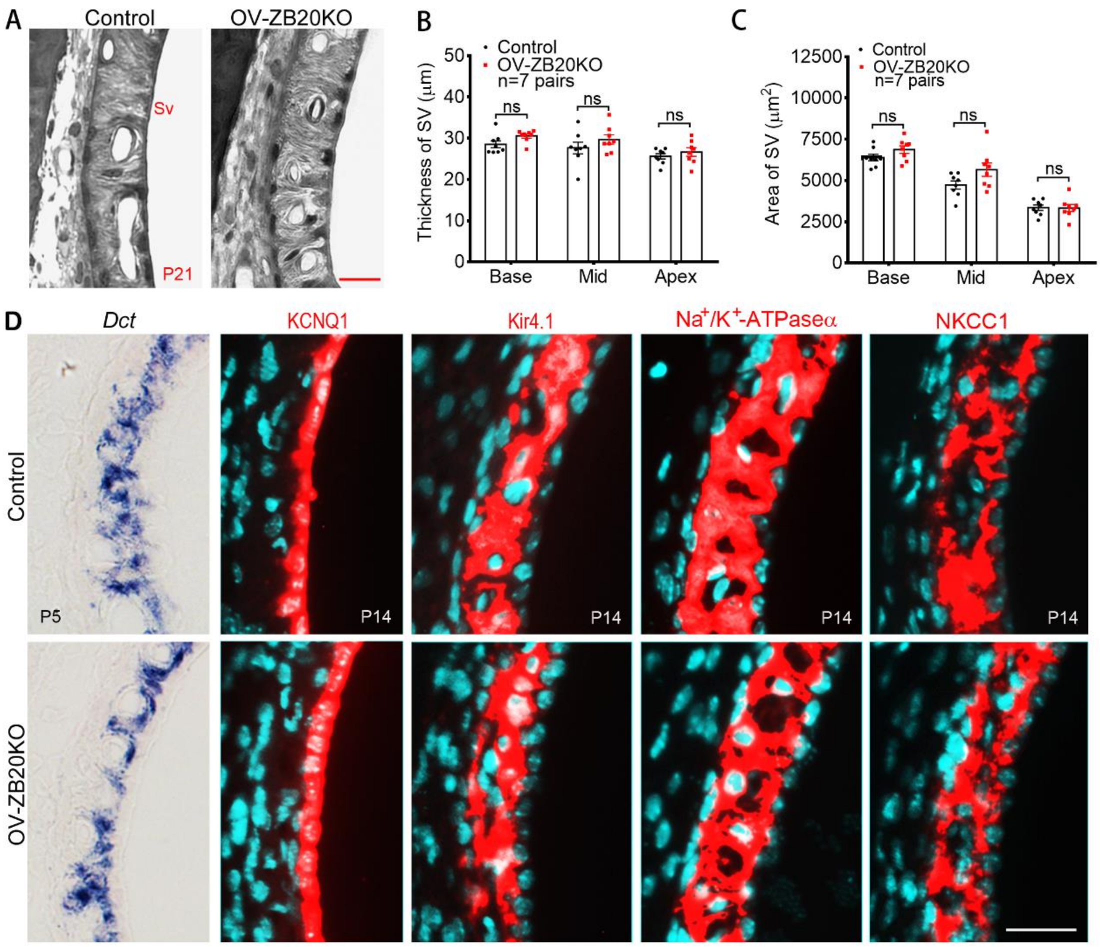


**Fig. S10. Ablation of *Zbtb20* does not affect the development of stria vascularis (SV).** (**A-C**) Representative semi-thin toluidine blue-stained cochlear SV sections showing comparable histology of SV between Control and OV-ZB20KO mice at P21. Graphs showing comparable thickness (**B**) and area (**C**) of SV between Control and OV-ZB20KO mice at P21. n=7 mice/group. Values are represented as mean ±SEM. A two-way ANOVA followed by post-hoc pairwise tests (Bonferroni method) was used to compare differences between the two groups. ns: not significant. (**D**) Representative cochlear SV sections at indicated ages showing comparable expressions of dopachrome tautomerase (*Dct*) mRNA (blue, revealed by in situ hybridization), KCNQ1, Kir4.1, Na+/K+-ATPase α and NKCC1 (red, revealed by immunostaining, nuclei were stained with DAPI in turquoise) between Control and OV-ZB20KO mice. n=5 mice/group for each molecular marker. Scale bars: 25 µm (**A**) and 50 µm (**D**).

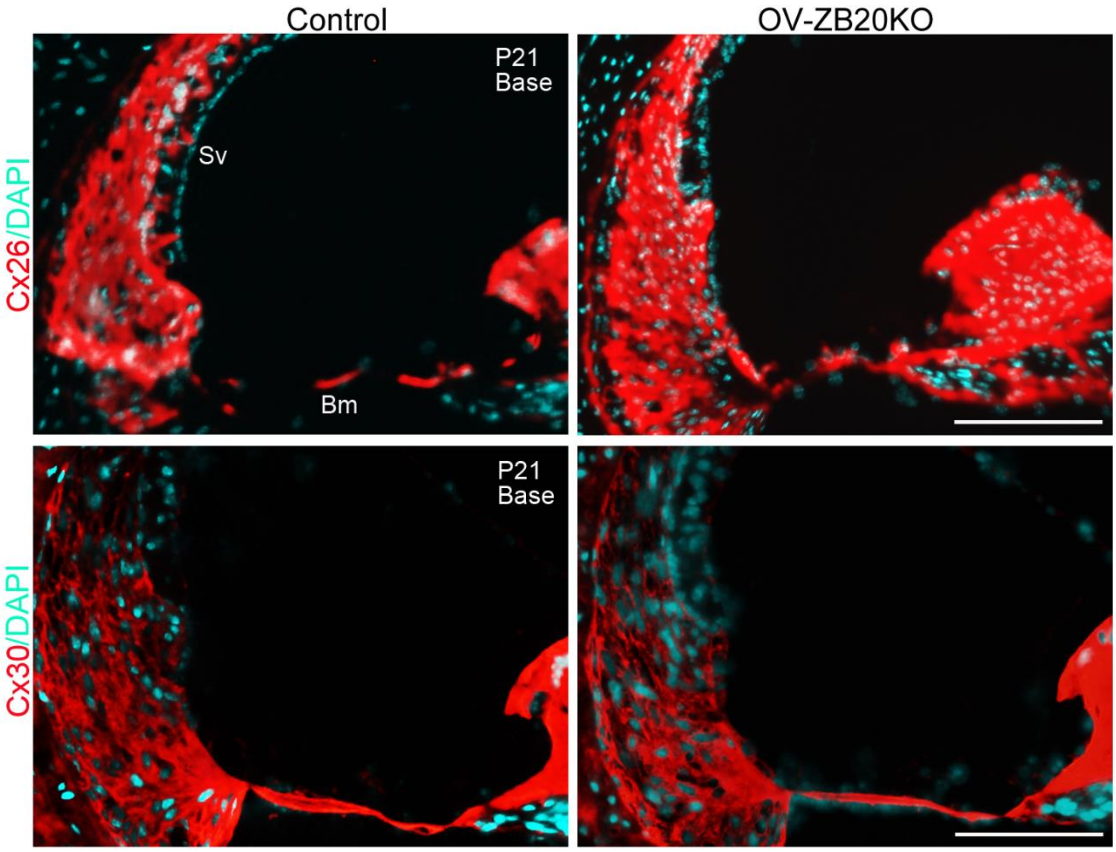


**Fig. S11. Ablation of *Zbtb20* does not affect the expression patterns of connexins in cochleae.** Representative immuno-stained cochlear sections showing comparable expression patterns of CX26 and CX30 (red) between Control and OV-ZB20KO mice at P21. Nuclei were stained with DAPI (turquoise). Bm, basilar membrane; Sv, stria vascularis. Scale bars: 100 μm. n=4 mice/group.

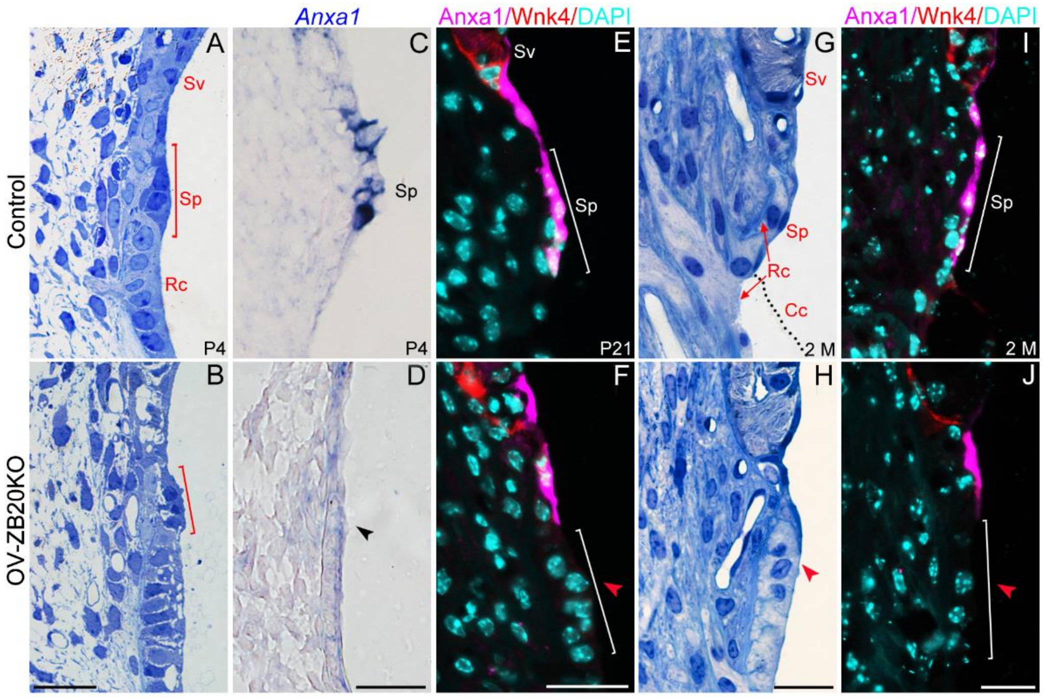


**Fig. S12. Ablation of *Zbtb20* disrupts postnatal development of spiral prominence. (A-B)** Representative semi-thin toluidine blue-stained cochlear sections from both control and OV-ZB20KO mice at P4 showing several spiral prominence (Sp) epithelial cells (more darkly stained, indicated by red bracket) located between the inferior tip of stria vascularis (Sv) and superiormost root cell (Rc). **(C-D)** In situ hybridization showing Anxa1 mRNA was detected in WT control but not OV-ZB20KO spiral prominence epithelium at P4. **(E-F)** Representative immuno-stained cochlear sections showing an absence of Anxa1 expression (magenta) in OV-ZB20KO SP epithelium at P21. SV was demarcated by Wnk4-positive (red) basal cells. Nuclei were stained with DAPI (turquoise). **(G-H)** Representative semi-thin cochlear sections showing a flattened SP in OV-ZB20KO mice at 2 months while WT counterparts exhibiting a mature morphology with squamoid epithelial cells directly connected with Claudius cells (Cc, indicated by black dotted line) covering the underlying root cells. **(I-J)** Representative immuno-stained cochlear sections showing an absence of Anxa1 expression (magenta) in OV-ZB20KO SP epithelium at 2 months. Scale bars: 20 µm in (A-B and G-H), 25 µm in(C-D and G-J). n=4 mice/group.

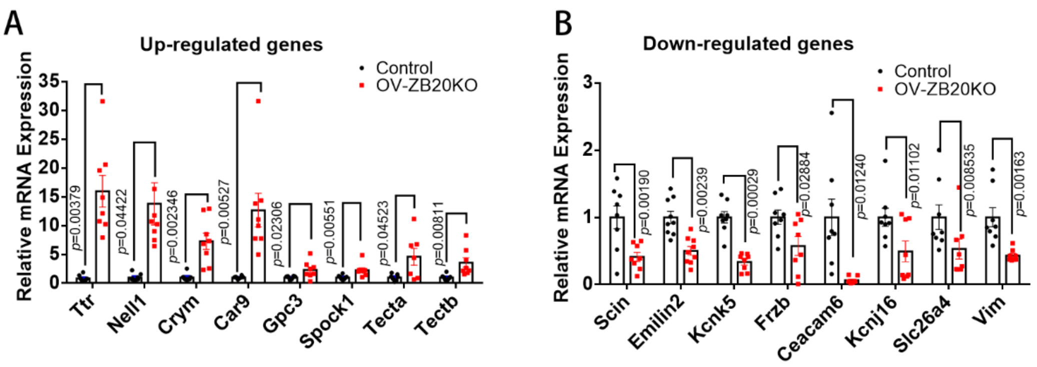


**Fig. S13. Validation of differentially expressed genes (DEGs) revealed by RNA-seq.** (**A-B**) Validation of DEGs revealed by RNA-seq using Real-time RT-PCR, showing relative mRNA expression levels (fold change) of up-regulated (**A**) and down-regulated (**B**) genes on cochlear epithelium from OV-ZB20KO mice and their WT controls at P10. n≧7 pairs for each group. Values are shown as means ±SEMs. Comparisons of two groups were analyzed using paired Student's t test (two-tailed). *p* values are indicated in figures.
